# Genetic mapping of flowering time and plant height in a maize Stiff Stalk MAGIC population

**DOI:** 10.1101/2022.01.31.478539

**Authors:** Kathryn J. Michel, Dayane C. Lima, Hope Hundley, Vasanth Singan, Yuko Yoshinaga, Chris Daum, Kerrie Barry, Karl W. Broman, C. Robin Buell, Natalia de Leon, Shawn M. Kaeppler

## Abstract

The Stiff Stalk heterotic pool is a foundation of US maize seed parent germplasm and has been heavily utilized by both public and private maize breeders since its inception in the 1930’s. Flowering time and plant height are critical characteristics for both inbred parents and their test crossed hybrid progeny. To study these traits, a six parent multiparent advanced generation intercross (MAGIC) population was developed including maize inbred lines B73, B84, PHB47 (B37 type), LH145 (B14 type), PHJ40 (novel early Stiff Stalk), and NKH8431 (B73/B14 type). A set of 779 doubled haploid lines were evaluated for flowering time and plant height in two field replicates in 2016 and 2017, and a subset of 689 and 561 doubled haploid lines were crossed to two testers, respectively, and evaluated as hybrids in two locations in 2018 and 2019 using an incomplete block design. Markers were derived from a Practical Haplotype Graph built from the founder whole genome assemblies and genotype-by-sequencing and exome capture-based sequencing of the population. Genetic mapping utilizing an update to R/qtl2 revealed differing profiles of significant loci for both traits between 636 of the DH lines and two sets of 571 and 472 derived hybrids. Genomic prediction was used to test the feasibility of predicting hybrid phenotypes based on the *per se* data. Predictive abilities were highest on direct models trained using the data they would predict (0.55 to 0.63), and indirect models trained using *per se* data to predict hybrid traits had slightly lower predictive abilities (0.49 to 0.55). Overall, this finding is consistent with the overlapping and non-overlapping significant QTL found within the *per se* and hybrid populations and suggests that selections for phenology traits can be made effectively on doubled haploid lines before hybrid data is available.

**Core Ideas:** A multi-parent advanced generation intercross (MAGIC) mapping population was developed from six founder Stiff Stalk maize inbreds with commercial relevance. Genetic mapping utilizing an update to R/qtl2 was demonstrated for flowering and plant height traits.

Genetic mapping using maize inbred and hybrid information was compared and provided insight into trait expression in inbreds relative to heterotic testcross hybrids.

## INTRODUCTION

Multi-parent mapping populations are an effective tool for discovering quantitative trait loci (QTL) in plant and animal species. Multi-parent advanced generation intercross (MAGIC) populations offer a powerful QTL mapping structure because intercrossing more than two parents increases genetic diversity while managing minor allele frequency and reducing haplotype length through recombination (reviewed in Scott et al., 2020). MAGIC populations have been used to successfully dissect the genetic control of complex traits in various plant species, including Arabidopsis (Kover et al. 2009), maize (Dell’Acqua et al. 2015), rice (Ogawa et al. 2018), barley (Sannemann et al. 2015), wheat (Gardner et al. 2016), sorghum (Ongom and Ejeta 2018), tomato (Pascual et al. 2015), and cowpea (Huynh et al. 2018). Multi-parent populations balance the advantages and disadvantages of biparental mapping populations and association panels. Geneticists often rely on the cross of two individuals with contrasting phenotypes to generate a population of segregating individuals and then perform linkage analysis to associate genetic loci with the trait of interest. Recently, increased marker density due to technological advancements and rapidly declining genotyping costs allowed researchers to evaluate diverse association panels to assay historical recombination to find associations between markers and phenotypes (reviewed in Tibbs Cortes *et al*. 2021). Despite the success of these methods, both techniques face intrinsic challenges. Biparental populations rely on the genetic diversity found in just two parents, which can limit the scope of discovered QTLs to the backgrounds studied. Association panels often contain rare alleles that do not meet the minor allele frequency threshold and are discarded due to low statistical power associated with such rarity. Thus, MAGIC populations seek to balance these characteristics by incorporating more than two genetic backgrounds while balancing minor allele frequency and increasing mapping resolution.

To study the genetic architecture of traits relevant to maize hybrid performance and adaptation, we developed a MAGIC population from six inbred lines spanning the diversity of the Stiff Stalk heterotic pool. The Stiff Stalk heterotic group was founded in the Iowa Stiff Stalk Synthetic (BSSS) breeding population, which was initiated during the 1930’s by Dr. George Sprague to improve stalk quality, yield, and agronomic quality of maize inbreds (Troyer 2004). Several key inbreds were released out of BSSS, including B14 in 1953, B37 in 1958, B73 in 1972, and B84 in 1979 (Russell 1972; Russell 1979; Troyer 1999). Since their release, these founder BSSS inbreds have been used extensively by breeders in the public and private sectors in the United States, and the Stiff Stalk group has become the *de facto* source of seed parent germplasm for many hybrid breeding programs. It is estimated that B73, B14, and B37 contributed conservatively 16.4% to germplasm released by Monsanto Company, Pioneer Hi-Bred, International, and Syngenta between 2004-2008 (Mikel 2011). In a group of 1,506 lines released under Plant Variety Protection (PVP) certificates between the year 2000 and 2019, researchers found that a third of the lines had kinship estimated Stiff Stalk admixture greater than 30%, and 15% of lines had Stiff Stalk admixture greater than 50% (White et al. 2020). Thus, the Stiff Stalk heterotic group remains a vital source of commercial maize germplasm in North America. This research utilized six Stiff Stalk inbreds - B73, B84, NKH8431, LH145, PHB47, and PHJ40 – that represent key founders in commercial breeding programs. Recent work reported the genome sequences of these inbreds (excluding B73) and found extensive genomic variation between B73 and the other five parents along with conservation of base BSSS haplotypes within each inbred (Bornowski et al. 2021).

Throughout the process of developing new inbred lines and hybrid varieties, maize breeders balance selecting for hybrid yield with other traits needed for successful inbred and hybrid seed production. Traits such as flowering time and plant height are vital to the success of an inbred within the breeding program and as a parent to a successful hybrid variety. Extensive research has been conducted on maize flowering time, including the discovery of a multitude of small to large effect QTL contributing to flowering time and photoperiod sensitivity variation in maize (Buckler et al. 2009; Xu et al. 2012; Wang et al. 2021) and the identification of several genes and regulatory elements involved in the pathway, including *ID1*, *DLF1*, *ZmCCA1*, *ZmMADS1*, *ZmCOL3*, *Vgt1*, *ZCN8*, *ZmCCT*, *ZmCCT9*, *ZmCCT10*, and *ZmMADS69* (Colasanti et al. 1998; Muszynski et al. 2006; Salvi et al. 2007; Wang et al. 2011; Hung et al. 2012; Alter et al. 2016; Jin et al. 2018; Huang et al. 2018; Guo et al. 2018; Liang et al. 2019; Stephenson et al. 2019). Flowering time and photoperiod sensitivity are determinants of maize yield because the combination leads to adaptation of maize lines to their intended environments, such that tropical lines with daylight sensitivity must undergo extensive selection for adaptation to succeed in northern regions that do not meet daylight needs of tropical plants (Xu et al. 2012). In addition, timing of flowering can influence the total length of time available for grain filling post flowering and the ability of a hybrid to mature within a frost-free seasonal interval. Within an environment, maize hybrids with full-season relative maturities often yield more than their shorter-season counterparts, and timing of planting date to achieve flowering time before environment specific cutoffs is vital for maximizing yield potential (Baum et al. 2019). However, later-maturing varieties can face risk due to early frosts and susceptibility to seasonal drought effects (Duvick and Cassman 1999), therefore plant breeders need to carefully balance flowering time and total maturity to suit their target population of environments to maximize maize grain yield. In maize hybrid varieties, flowering time typically exhibits heterosis where the hybrid flowers sooner than the earlier of the two inbred parents, as demonstrated by a partial diallel of ex-PVP inbreds (Li et al. 2018) and an association panel of 302 diverse inbreds crossed to B73 (Flint-Garcia et al. 2009).

Like flowering time, substantial efforts have been devoted to understanding the genetic underpinnings of maize plant and ear height. Major mutations in the gibberellin and brassinosteroid pathways affecting height have been identified in addition to numerous QTL (reviewed by Salas Fernandez et al., 2009). Despite its high heritability, QTL affecting height tend to have very small effects, with the largest effect in the maize US-NAM population explaining 2.1 +- 0.9% of the variation, which suggests that maize height follows an infinitesimal model of inheritance (Peiffer et al. 2014). In addition, identification of QTL can depend on environmental conditions such as drought and nutrient stress, which may reduce the relative proportion of additive genetic variance compared to genotype by environment and error variance (Cai et al. 2012; Wallace et al. 2016). In general, taller plants can face increased root and stalk lodging pressure due to the proportionally higher placement of the ear on the stalk, which increases the ear’s leverage during wind events or disease pressure. During the Green Revolution, major yield gains were made in rice and wheat by decreasing overall plant height, which reduced the risk of lodging under more intensive agricultural management (reviewed by Khush, 2001). Maize breeders consider height selection in both inbreds and hybrids, as lodging can make harvest difficult and inefficient for both seed parents and commercial varieties. Due to heterosis, the hybrid is usually taller than the taller of the two inbred parents, as shown in a partial diallel of ex-PVP inbreds (Li et al. 2018) and in an association panel of 302 diverse lines crossed to B73 (Flint-Garcia et al. 2009).

The main objectives of this work are to: i) report a MAGIC population based on the Stiff Stalk heterotic group and its associated genetic and phenotypic resources, ii) dissect the genetic architecture of flowering time and plant height within the *per se* population of DH lines and two test cross populations, and iii) perform genomic prediction to investigate the relationship between *per se* and hybrid phenotypes.

## MATERIALS AND METHODS

### Population Development

Inbreds B73, B84, NKH8431, LH145, PHB47, and PHJ40 were chosen to represent the primary Stiff Stalk sub-heterotic groups (Table 1) (White et al. 2020). Biographical information for each line was obtained from the Germplasm Resource Information Network (GRIN) database (npgsweb.ars-grin.gov). The inbreds B73 and B84 were released from the BSSS in cycles five and seven, respectively, and B84 contains resistance to *Helminthosporium turcicum* (“Ht” currently known as *Setosphaeria turcica*, common name Northern Corn Leaf Blight). Inbred LH145 was developed by Holden’s Foundation Seed, Inc. (acquired by Monsanto Company in 1997) from the cross of A632Ht and CM105, both of which have B14 as a parent. Inbred NKH8431 was developed from one B73 derived line and two B14 derived lines by Northrup, King & Company. Inbreds PHB47 and PHJ40 were both released by Pioneer Hi-Bred, International (PHI). Inbred PHB47 was made from a cross between B37 and SD105, with two backcrosses to B37 before inbred development. Inbred PHJ40 is an early flowering flint and Stiff Stalk line developed in Ontario, Canada, with previously demonstrated admixture with B37 (White et al., 2020). All inbred lines except B73 previously underwent whole genome, reference guided assembly, which revealed extensive genetic and genomic diversity between the five lines and B73 (Bornowski et al. 2021).

**Table 1.** Origins of Stiff Stalk inbred lines

Sub-heterotic groupings from White et al., 2020
| Line | Originator | Sub-heterotic group | PI number |
| --- | --- | --- | --- |
| B73 | Iowa State University | B73 | PI 550473 |
| B84 | Iowa State University | B73 | PI 608767 |
| LH145 | Holden's Foundation Seed, Inc. | B14 | PI 600959 |
| NKH8431 (alias H8431, NPH8431) | Northrup, King & Company | B14 | PI 601610 |
| PHB47 (alias B47) | Pioneer Hi-Bred International, Inc. | B37 | PI 601009 |
| PHJ40 | Pioneer Hi-Bred International, Inc. | Flint | PI 601321 |

The population, named WI-SS-MAGIC, was initiated at the University of Wisconsin during summer 2008. The six parents were crossed in a half diallel. Next, every possible F_1_ hybrid combination cross was attempted, and seed was included in the subsequent balanced bulk to maintain equal representation of all parents and account for any failed crosses. In subsequent generations, plants from the population bulk were randomly intermated by designating each plant as a pollen parent or seed parent and using the individual only once for crossing. Balanced bulks were made after harvest of the first intermating and a subset of the population (hereafter “Subset A”) was sent for doubled haploid (DH) induction, provided as in-kind support by AgReliant Genetics. The remaining balanced bulk was randomly intermated for two additional generations and then sent for DH induction (hereafter “Subset B”). Individuals in Subset A were given coded names beginning with W10004 and numbered from 1 to N, where N is the number of individuals (i.e. W10004_0001 through W10004_04xx), and individuals in Subset B were named using W10004 and a number from 500 to 500+N, where N is the number of individuals returned (i.e. W10004_0500 through W10004_xxxx) (Table S1).

### Collection of *Per Se* DH Line Phenotypic Data

A set of 779 DH lines was planted during summers 2016 and 2017 at the West Madison Agricultural Research Station in Verona, WI (Table S1). Subset A and Subset B groups were organized as subblocks within a randomized complete block (RCB) design with two replications. Parents were included as checks in both subblocks. Both trials were planted in fields that followed soybeans in the previous year and were managed with standard agronomic practices. Detailed information about planting dates and densities, soil types, and nutrient and pesticide management is presented in Supplemental Table 2. Three representative plants per plot were measured for plant and ear height. Plant height was measured as the height from the ground, in centimeters, to the collar of the flag leaf, while ear height was the height, in centimeters, from the ground to the base of the node subtending the uppermost ear. Growing degree units to anthesis and silking were measured on a whole plot basis (AnthGDU and SilkGDU, respectively). Anthesis and silking were measured as the number of days from planting it took to observe approximately 50% of the plants in the plot to reach pollen shed and silk extrusion, respectively. Dates were converted to growing degree units using a base temperature of 50° F and maximum temperature of 86° F (Pope 2008) using temperature data obtained from the weather station located at the University of Wisconsin (UW) West Madison Agricultural Research Station to standardize for differential daily heat unit accumulation across years. Since lines developed through doubled haploidy are expected to be genetically uniform, lines with observable phenotypic segregation were discarded. Severely lodged plants were not evaluated for height characteristics. To remove outlier data points, individual plant measurements were discarded if the ear height to plant height measurement ratio was less than 0.25 or greater than 0.75, and whole plot ear or plant height measurements were discarded if the within plot variance was greater than 500 cm^2^.

### Generation of Hybrids and Collection of Phenotypic Data

Hybrid seed was produced by crossing the WI-SS-MAGIC population to PHJ89 and DKH3IIH6 (hereafter 3IIH6). The hybrid populations were named SS-PHJ89 and SS-3IIH6. PHJ89 is an Oh43-type inbred line developed by Pioneer Hi-Bred (White et al. 2020). The inbred 3IIH6 is an Iodent-type inbred line developed by DeKalb Genetics Corporation (acquired by Monsanto in 1998, now owned by Bayer AG) through selfing the F_1_ Hybrid PHI3737 (Dekalb Plant Genetics 1994). PHJ89 and 3IIH6 are related by pedigree through their founder PHG47, which is one of the two parents of PHJ89 and one of the parents of hybrid variety PH3737 from which 3IIH6 was generated through selfing, so they are expected to contain regions of identity by descent (Pioneer Hi-Bred International Inc. 1992; Mikel 2011). Hybrids were grown during summers 2018 and 2019 at the UW West Madison Agricultural Research Station in Verona, Wisconsin and at the UW Arlington Research Station in Arlington, WI. Hybrids were blocked by tester, and each block included at least five replicates each of two commercial hybrids (DKC50-08RIB and DKC54-38RIB) and two replicates of each respective population parent-tester combination, when seed was available. All trials were incompletely replicated, where each hybrid genotype was grown at least once in each experiment with a consistent random subset replicated a second time. A total of 689 SS-3IIH6 hybrids were grown, of which 316 were replicated, while a total of 561 SS-PHJ89 hybrids were grown, of which 377 were replicated (Table S1). The same set of replicated and unreplicated lines were grown across years and locations, with unique plot randomizations for each year-location combination. Replicated hybrids were randomized among the unreplicated hybrids within their respective tester blocks. All trials were planted in fields that followed soybeans in the previous year and were managed with standard agronomic practices.

Detailed information about planting dates and densities, soil types, and nutrient and pesticide management is presented in Table S2. Flowering time growing degree units were recorded in the same manner as previously described for the *per se* population using weather data obtained from each research station. Flowering time was recorded for all hybrid plots in West Madison and for approximately 36% and 33% percent of hybrid plots in Arlington in 2018 and 2019, respectively. Plant height and ear height were measured on three competitive plants per plot for all plots. Stand counts were recorded manually, and plots were discarded if they contained fewer than twenty plants. The 2019 West Madison trial experienced extremely wet and cold germination conditions, which led to low stand counts for the SS-3IIH6 block. Only 19% and 38% of the plots had stands greater than 50 and 40 plants, respectively, which prompted us to discard the height data due to inconsistent interplant competition but keep flowering time data due to good correlations with flowering time from the previous year. To remove outlier data points, individual plant measurements were discarded if the ear height to plant height measurement ratio was less than 0.25 or greater than 0.75, and whole plot height measurements were discarded if the within plot variance was greater than 500 cm^2^. Plot average height measurements and flowering GDU measurements that were more than three standard deviations away from the experiment wide mean were discarded.

### Analysis of Phenotypic Data

A two stage approach was taken to analyze plot mean phenotypic data (Table S3). In stage one, following the procedures of (Rogers et al. 2021), linear mixed models were fit using R/ASReml version four (Butler et al. 2017; RCoreTeam 2018) for each population within each environment using genotype as a fixed effect and replicate as a random effect. Next, models were fit with all combinations of autoregressive first order (AR1) and IID residual covariance structures of the x and y grid coordinates of the plot locations to account for spatial variation. The model with the lowest Schwarz’s Bayesian Information Criterion (BIC) (Schwarz 1978) was chosen to represent the environment, and the BLUEs and standard errors were extracted from the model (Table S4). Due to our incomplete block structure, the residual spatial correlations were restricted to -0.6 < r < 0.6. Any models with correlation outside this range were reset to using no residual covariance structure. In stage two, genotypes were fit as fixed effects and environment and genotype by environment interaction terms were set as random effects. To weight the second stage analysis, the reciprocals of the first stage BLUE standard errors were carried forward, which represent the genotype by replication interactions, and the units variance was constrained to one. Within each experiment, any phenotypic BLUE that fell outside 2.5 times the interquartile range (IQR) was discarded as an outlier. Following the data cleaning described in the previous sections and the *post hoc* cleaning based on IQR, BLUEs were calculated for 730 DH lines, 658 SS-3IIH6 hybrids, and 535 SS-PHJ89 hybrids. Parental check lines were included in the analysis because they constitute the same population as the experimental lines, while commercial check hybrids were not included in the analysis. To estimate variance components and calculate heritability, the same model was used except genotype was set as a random effect. Heritability was calculated as follows (Cullis et al. 2006):

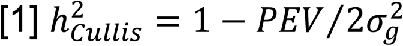

using the prediction error variance (PEV) and genetic variance 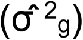 obtained from the stage two analysis. To compare phenotypic variances across populations, the squared coefficient of variation was calculated to correct for the differences in scale between *per se* and hybrid phenotypes (Falconer and Mackay 1989). Pearson correlations within and between phenotypes were calculated on trait BLUE values within and between the DH and two hybrid populations.

## Genetic Data

### Sequencing Using Exome Capture

Exome capture sequencing was performed on 701 DH lines from the WI-SS-MAGIC (Table S1) using a custom capture design acquired from Roche Diagnostics Corporation (Indianapolis, IN). Probes were designed to target the 5’ and 3’ ends of the untranslated regions (UTRs) of the maize B73_RefGen_v2 genic regions and presence absence variation (PAV) regions derived from alignment of whole genome sequencing reads of a core set of 32 inbreds to B73 (Brohammer et al. 2018; Mazaheri et al. 2019). In total, 82,351 genic regions (approximately 26.5 Mb) and 492 PAV regions (approximately 2.8 Mb) of the maize genome were targeted using tiled, variable length probes, with an average probe size of 75 nt (File S1). Any overlapping regions were collapsed into a single target. The target regions ranged in size from 50 to 49,777 nucleotides (nt), with a mean size of 353.6 nt (File S2). Briefly, DNA was extracted using seedling tissue using a modified CTAB method (Saghai-Maroof et al. 1984), sheared, and hybridized with adapters prior to SeqCap EZ solution capture, as previously described (Mascher et al. 2013) (File S3).

DNA was then amplified, enrichment quality control performed, and libraries sequenced by the United States Department of Energy Joint Genome Institute (JGI) in paired end mode on the Illumina HiSeq 2500. Raw sequence quality was evaluated using FastQC v0.11.5 (https://www.bioinformatics.babraham.ac.uk/projects/fastqc) and MultiQC v1.0 (Ewels et al. 2016). Reads were then trimmed, low quality bases removed, and adapters removed using Cutadapt v1.14 (Martin 2011) with the following parameters: -- length 150, -m 20, -q 20, 20, --times 2, and -g/-a/-G/-A along with their respective adapter sequences. After cleaning, read quality and adapter content were evaluated again using FastQC v.0.11.5 and MultiQC v1.0.

### Genotyping By Sequencing

Additional genotyping was performed on 144 DH lines using Genotyping-by-Sequencing (GBS) at the University of Wisconsin Biotechnology Center (Table S1). Briefly, dual digest GBS was performed with restriction enzymes *Pst*I and *Msp*I on frozen seedling leaf tissue (Elshire et al. 2011; Poland et al. 2012). DNA was sequenced using an Illumina NovaSeq6000 in paired end mode 150 nt and analyzed using bcl2fastq v2.20.0.422 (San Diego, CA, USA). Read one was demultiplexed and barcodes were removed using Tassel-5-Standalone plugin “ConvertOldFastqToModernFormatPlugin” with parameters “-e PstI-MspI” and “-p R1” (Bradbury et al. 2007). Read two was not included in future analysis.

### Practical Haplotype Graph

A Practical Haplotype Graph (version 0.0.30) was built using B73 v5 as the reference (Bradbury *et al*. 2021, maizegdb.org). The B73 RefGen_v5 annotation of genes (Zm-B73-REFERENCE-NAM-5.0_Zm00001eb.1.gff3, available at maizegdb.org) was used to make the initial reference ranges, which were supplied to the “CreateValidIntervalsFilePlugin” to collapse any overlapping ranges and format for input into PHG. B73 RefGen_v5 was loaded as the reference assembly using the “MakeInitialPHGDBPipelinePlugin”, followed by the other five parental *de novo* genome assemblies using the “AssemblyHaplotypesMultiThreadPlugin” (Bornowski et al. 2021). The “AssemblyHaplotypesMultiThreadPlugin” uses mummer4 to align each assembly to the reference by chromosome in parallel (Marçais et al. 2018). Next, B73 was added to the graph using the “AddRefRangeAsAssemblyPlugin” which allows B73 haplotypes to be included as potential parental sequences.

A ranking file was generated by counting the number of haplotypes found within each assembly. The ranking file is necessary when two or more haplotypes are collapsed into a consensus haplotype, where the sequence of the highest-ranking line will represent the group. Consensus haplotypes were made using the PHG shell script “CreateConsensi.sh” with parameters “mxDiv=0.0001” and “minTaxa=1”. All other parameters were left as default. The value of “mxDiv=0.0001” was chosen such that genic regions would only be collapsed if they were truly identical, since over 90% of maize gene models are shorter than 10,000 bp. After consensus haplotypes were generated, the “ImputePipelinePlugin” with parameter “-imputeTarget pathToVCF" was used to index the pangenome, map exome capture and GBS reads to the graph, use a Hidden Markov Model to find paths through the graph for each taxon, and call SNPs in the genic reference regions for the progeny population. Exome capture reads were aligned as paired end sequences, while the GBS reads were aligned as single end sequences. Parental assembly genic SNPs were extracted from the graph using the “FilterGraphPlugin” and “PathsToVCFPlugin”. Due to the expected homozygosity of the DH lines and parental assemblies, only homozygous SNPs were generated from the PHG.

Markers were filtered and selected for mapping using Tassel-5-Standalone (Bradbury et al. 2007). The combined file of parental and population individuals (File S4) was filtered to remove any non-Stiff Stalk individuals that were included as checks, SNPs with any missing parental data were removed, minor SNP states were set to missing to remove third, fourth and other alleles, and the SNP was removed if the minor allele frequency was less than 0.05. To reduce correlation between SNPs and decrease QTL mapping computational time, 100,000 evenly spaced SNPs were selected across the ten chromosomes and converted to numerical major or minor alleles.

### Population genetic characteristics

Multidimensional scaling (MDS) was performed using 1.8 million unfiltered genic SNPs to confirm lack of population structure. A distance matrix was calculated using the “DistanceMatrixPlugin” from Tassel-5- Standalone with default parameters (Bradbury et al. 2007). In R, cmdscale() was used to calculate classical MDS on the distance matrix for the first two dimensions (RCoreTeam 2018). Allele frequencies were calculated on the set of 100,000 SNPs used for QTL mapping.

### QTL Mapping

Quality control analyses, single-marker QTL mapping, and SNP associations were performed using R/qtl2 with the cross-type corresponding to our mating design, “dh6” (Broman et al. 2019). Whenever individuals underwent both exome capture and GBS, the GBS reads were used to generate markers for QTL mapping. To prepare the data, a control file was created using the function write_control_file() from R/qtl2, which specifies the cross type for our population, the file names of the population and parental SNPs, the physical map coordinates for the SNPs, the phenotype file, the cross information file, which contains the number of meiosis used to generate each individual, and the parental alleles “AA” through “FF”.

Due to the high density of markers, a genetic map was approximated by converting each SNP’s megabase pair position to centiMorgans using the B73 RefGen_v5 chromosomal genome length of 2132 Mbp divided by the composite US-NAM genetic map length of 1456.68 cM (Li et al. 2015). The control file and all materials needed for mapping are provided as Supplemental File S5.

Any line with segregating *per se* phenotypes had previously been removed from further analysis. To identify potential sample duplicates, the function compare_geno() was used to calculate marker matching for all pairwise comparisons, and any pair of individuals with greater than 95% marker sharing was removed. Conditional genotype probabilities, or the true genotype underlying the observed markers, were calculated using a Hidden Markov Model (HMM) in the function calc_genoprob(), with an error probability of 0.01. (Broman and Sen 2009, Appendix D). After calculating genotype probabilities, the maximum marginal probability of the parental haplotypes was calculated and the total number of crossover events per individual was identified using the function count_xo(). Crossover locations were estimated using the function locate_xo(). Lines with unusually high numbers of total crossovers could be the result of sample contamination during population development or maintenance, as the HMM cannot accurately deduce the correct underlying parental haplotypes in non-parental regions, and instead frequently switches back and forth among the parent haplotypes. Lines in Subset A with more than 150 crossovers or lines in Subset B with more than 250 crossovers were removed from further analysis.

After quality control, 657 individuals remained with phenotypes and genotypes for mapping purposes (Table S1). The genotype probabilities were used to calculate a kinship matrix, so the analysis could account for the relatedness between individuals using the “leave one chromosome out” (LOCO) method, which uses a kinship matrix derived from all other chromosomes except the chromosome under study to allow for a random polygenic effect (Yang et al. 2014). Next, single marker analysis was performed using a linear mixed model with the kinship matrix as a covariate to find associations between genotype and phenotype. Log of odds (LOD) thresholds were determined as the 95th percentile LOD score after 1,000 permutations of the founder probabilities using the function scan1perm() (Churchill and Doerge 1994; Cheng and Palmer 2013). Bayesian credible intervals for QTL peaks were calculated using the function find_peaks(), with LOD thresholds specific to each phenotype and probability of 0.95. To declare two QTL under one large peak, the LOD threshold was required to drop by at least five. Chromosome-wide QTL BLUP effects were calculated using the function scan1blup(), and single locus BLUP effects were estimated using fit1() with “blup=T”.

In addition to single marker QTL mapping, we also performed SNP association using the R/qtl2 function scan1snps(), with the same kinship matrix as previously described provided to account for population structure. Finally, not all DH lines included in the *per se* mapping were used to make the hybrid populations. To account for this difference in sampling between the *per se* traits and the hybrid traits, the sets of DH lines included in each hybrid population were used to repeat the mapping and permutation procedures for each *per se* trait that corresponded to a hybrid trait.

### Genomic Prediction

To test the correlation between *per se* and hybrid phenotype based on the DH population *per se* genetics, we performed genomic prediction using the 100,000 SNP markers used for QTL mapping. We used the package R/rrBLUP to perform ridge regression on the marker effects, which is equivalent to calculating genomic estimated breeding values using a realized relationship matrix (Hayes et al. 2009; Endelman 2011). We used fivefold cross validation to train and test the models predicting *per se* and hybrid traits. We partitioned the phenotypic data into five segments and used four segments for training the model and the remaining portion for testing the model. We predicted the phenotypes for each of the five testing segments and calculated the correlation between the predicted and observed phenotypes, which comprised one replication. We repeated this process 100 times for each of the phenotypes. To indirectly predict the hybrid phenotypes from the parental *per se* phenotypes, we calculated the correlation between the testing set predicted *per se* phenotypes and the observed hybrid phenotypes.

## RESULTS

### Phenotypic variation

Plant height and AnthGDU in the *per se* and hybrid populations showed normal distributions, and heritability ranged from 0.83 for SS-3IIH6 AnthGDU to 0.89 for SS-PHJ89 AnthGDU (Figure 1, Table S5). The genetic variance for *per se*, SS-3IIH6, and SS-PHJ89 AnthGDU was 2781.6 GDU^2^, 1148.6 GDU^2^, and 944.3 GDU^2^, respectively, while the genetic variance for *per se*, SS-3IIH6, and SS-PHJ89 plant height was 386.3 cm^2^, 150.1 cm^2^, and 144.6 cm^2^, respectively. Similarly, the squared coefficient of variation for *per se*, SS-3IIH6, and SS-PHJ89 AnthGDU was 18.00, 9.24, and 7.33, while the squared coefficient of variation for *per se*, SS-3IIH6, and SS-PHJ89 PH was 124.93, 26.79, and 22.35, respectively. All traits except *per se* and SS-PHJ89 AnthGDU exhibit transgressive segregation, where one or more progeny DH lines have more extreme values than all the parents (Figure 1). The parental line PHJ40 was the earliest flowering individual in the *per se* and SS-PHJ89 experiment.

**Figure 1:**
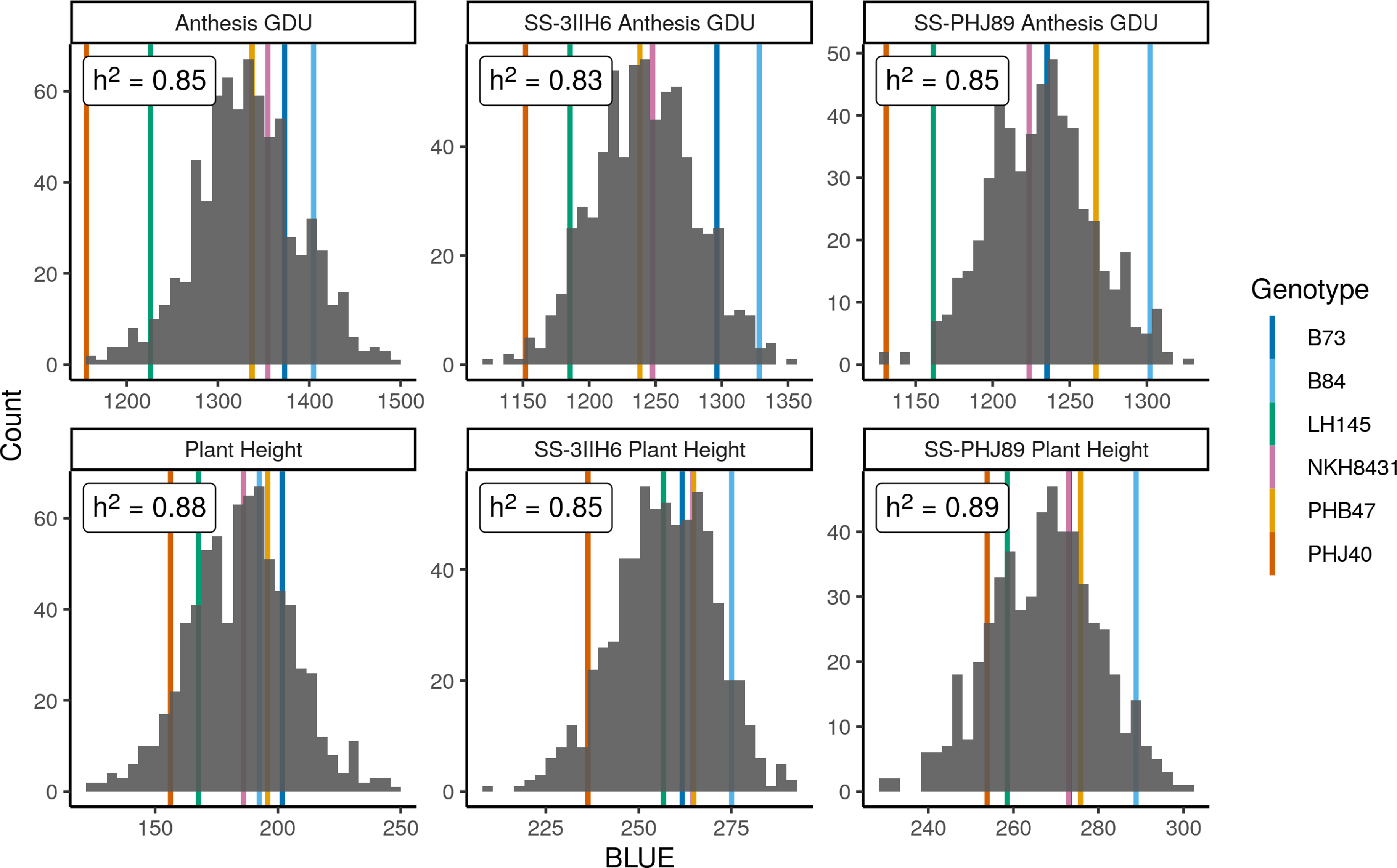
Distributions of phenotypic BLUEs and heritabilities Distributions for anthesis growing degree units (GDU) and plant height for the UW-MAGIC-SS *per se* population, SS-3IIH6 hybrid population, and SS-PHJ89 hybrid population. Trait heritabilities are in the upper left of each plot. Population parent BLUEs are plotted as colored lines behind each distribution.

Anthesis and silking are highly correlated within both DH lines and hybrids, ranging between Pearson r=0.83 for *per se* SilkGDU to *per se* AnthGDU to r=0.93 for SS-3IIH6 SilkGDU to AnthGDU (data not shown). The correlations between *per se* AnthGDU and SS-3IIH6 AnthGDU is r=0.64 and *per se* AnthGDU to SS-PHJ89 AnthGDU is r= 0.66 (Figure 2A and 2B). Correlation between AnthGDU for the two hybrid populations is higher at r=0.69 (Figure 2C). Plant height and ear height are also highly correlated within DH lines and hybrids. Correlations between plant height and ear height are r=0.79 within both the *per se* and SS-3IIH6 populations and r=0.83 within the SS-PHJ89 population (data not shown). *Per se* to hybrid plant height correlations are r=0.64 between DH lines and SS-3IIH6 and r=0.71 between DH lines and SS-PHJ89 (Figure 2D and 2E). Hybrid to hybrid plant height correlation is 0.72 (Figure 2F). The high correlation between hybrids is expected, due to both the highly additive nature of flowering time and height and the relatedness between testers 3IIH6 and PHJ89.

**Figure 2:**
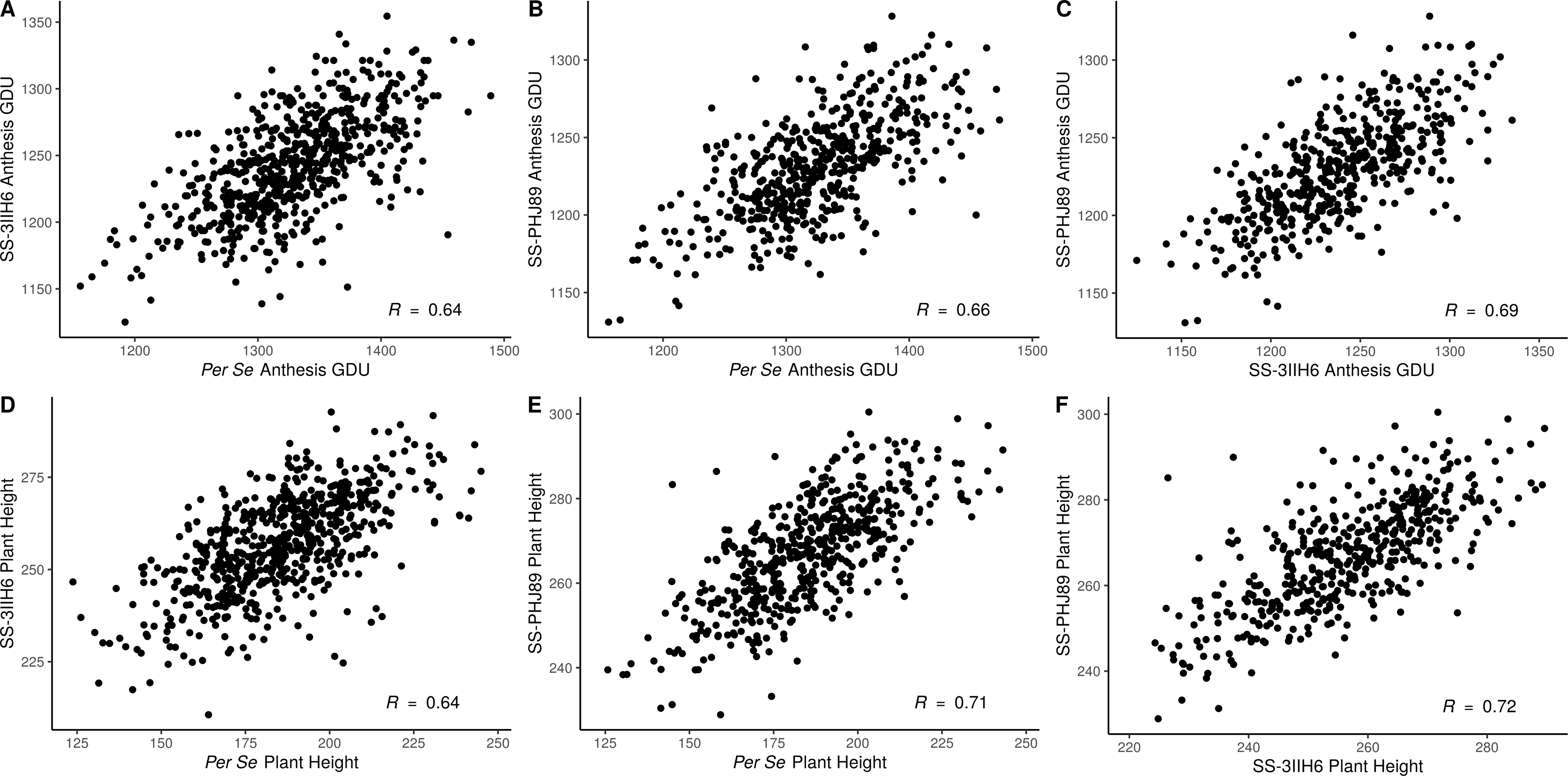
Phenotypic correlations between populations Scatterplots of BLUEs demonstrate the positive correlation within traits, between populations. Pearson correlations are shown in the lower right.

On average, the SS-3IIH6 population is 71.5 cm taller and sheds pollen 90.2 GDU earlier than its DH counterparts, and the SS-PHJ89 population is 82.0 cm taller and sheds pollen 101.1 GDU earlier than its DH founders. Finally, height and flowering are also correlated within populations, where r=0.35, 0.41, and 0.59 for the *per se,* SS-3IIH6, and SS-PHJ89 populations, respectively (Figure S1).

### Practical Haplotype Graph

The 39,035 B73 RefGen_v5 annotated gene models were used as initial reference ranges, and after collapsing overlapping ranges, 36,399 genic ranges remained, and 36,401 intergenic ranges were inserted between genic ranges for a total of 72,800 ranges. The average genic range width was 4755 bp, while the average intergenic range width was 53,811 bp. The theoretical maximum number of haplotypes per reference range is six, which represents either all inbreds having sequence that aligns to the reference (including reference to reference alignment), or five inbreds that have alignment with the reference and one with a missing haplotype. Not all assemblies have sequence that aligns to every range owing to structural variation between the parental genomes. Each parental assembly had different numbers of total haplotypes aligning to B73, from 51,070 haplotypes for PHJ40 to 66,890 for B84. PHJ40 is known to be more structurally diverse from the other founders (Bornowski et al. 2021), and the next lowest number of aligned haplotypes was 62,144 for LH145. After collapsing haplotypes into consensus sequences, the average number of haplotypes per PHG reference range was reduced from 5.7 to 3.3 (Figure 3A).

**Figure 3:**
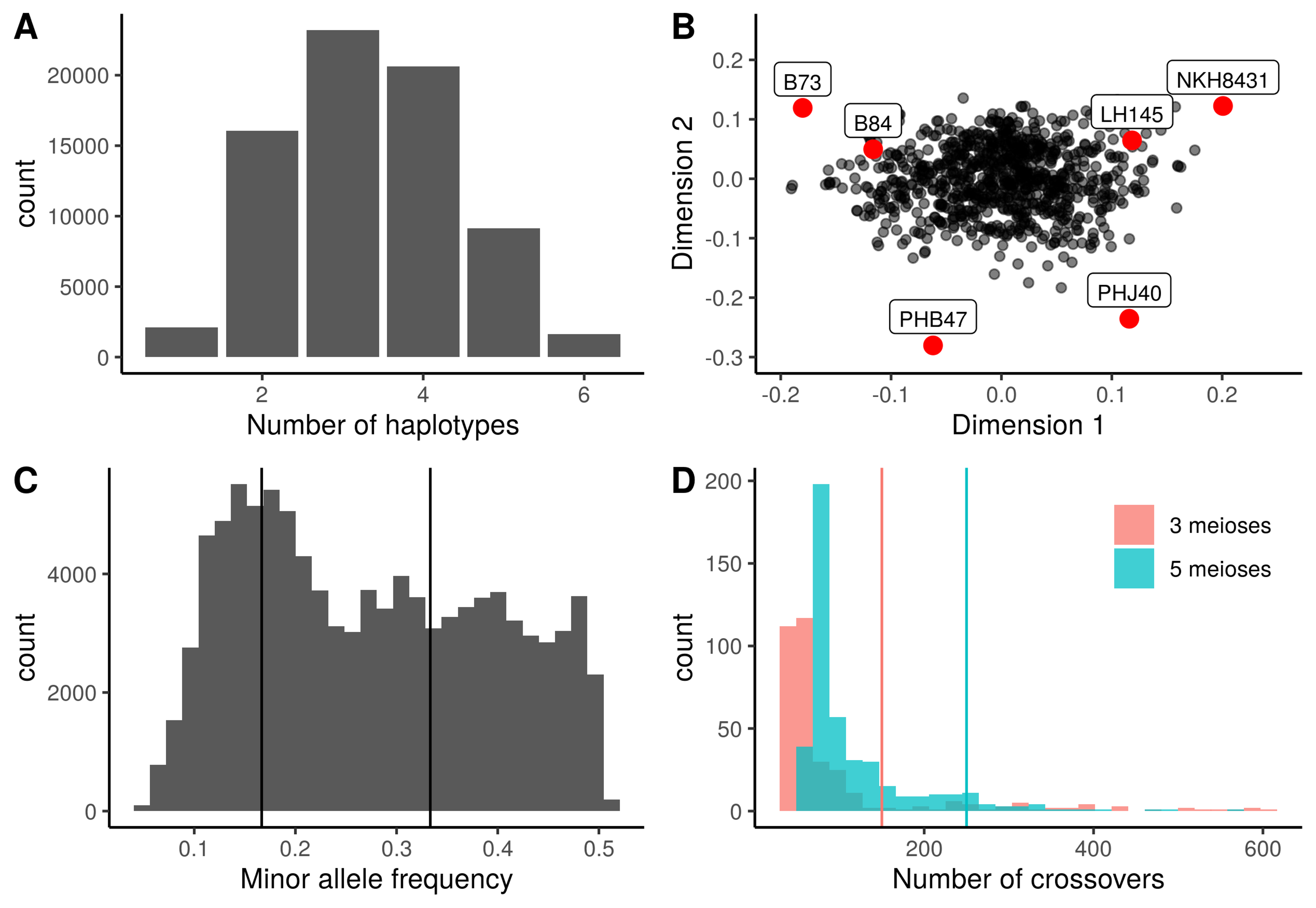
Population genetic characteristics (A) Distribution of the number of consensus haplotypes found per reference range. (B) Multidimensional scale plot using 1.8 million genic SNPs, with the population parents plotted in red. (C) Distribution of the minor allele frequencies for 100,000 filtered numeric SNPs, with vertical lines plotted at expected peaks of 1/6 and 2/6. (D) Histogram of the number of crossovers per individual for the two population subsets prior to filtering lines with high total crossovers. Six lines with more than 600 crossovers are not included. Vertical lines indicate the thresholds used for discarding lines at 150 crossovers for subset A (generated using three meioses) and 250 crossovers for subset B (generated using five meioses).

Identity by descent relationships are present between all of the lines due to their common heritage from the BSSS, and these relationships are strongest between B73 and B84, B73 and NKH8431, and NKH8431 and LH145. Consensus parental and population haplotype identification numbers are presented in Table S6.

### Population genetic characteristics

Multidimensional scaling (MDS) confirmed the lack of population structure within our population (Figure 3B). The parental inbred lines fall on the perimeter of the point cloud, with no discernable clustering of progeny individuals. Allele frequency distributions for the major and minor alleles appear as expected, with peaks near 1/6 and 2/6, corresponding to private and two-way sharing of alleles within the parents, respectively (Figure 3C). The founder probabilities and the total number of observable crossovers were calculated using R/qtl2. The two population subsets have overlapping distributions for the total number of crossovers per individual. While examining the locations and total numbers of crossovers present within individuals, we found some areas of the genome in certain individuals contained unusually high numbers of crossovers. Such areas indicate that the Hidden Markov Model fails to choose a single founder for the area, and instead rapidly switches between founders. While some individuals had high total genome wide incidence of crossovers, which indicates a sample mix-up, some lines had isolated areas of high crossover in only a few regions. Small areas of high crossover could be caused by several factors, including residual heterozygosity in the founder inbreds, introgression from the DH inducer (Li et al. 2009), contamination during population development from an inbred closely related to one of the founders, or technical issues during the SNP calling pipeline. In addition, using a PHG with imputation to generate SNPs for the population forces each individual to have haplotypes only from the population founders which complicates identifying areas of inducer introgression or contamination. Most importantly, QTL mapping results did not change significantly between the raw, full set of lines and the cleaned, reduced set of lines filtered for high total crossovers (150 crossovers for Subset A, 250 crossovers for Subset B), indicating that our results are robust to this low level of uncertainty.

After removing individuals with high numbers of total crossovers, the Subset A (three total meioses to generate DH lines) has an average of 60 crossovers, while Subset B (five total meioses to generate DH lines) has an average of 101 crossovers (Figure 3D). The parental haplotypes for a set of eight population individuals reveal a mosaic of the founder genotypes (Figure S2). The top row of individuals belongs to the Subset A and show longer parental haplotypes than the bottom row of individuals, which belong to Subset B. In many individuals, there are chromosome sections plotted in white, which correspond to areas where the founder probabilities do not rise above 0.5. This is expected, due to the related nature of the population founders and the segments of identity by descent between them. For example, large stretches of identity by descent between B73 and B84 due to their selection out of the BSSS would make assigning population haplotypes to either of the parents difficult, and this issue is compounded by the presence of BSSS lines in the pedigrees of the other population parents. After removing individuals with high crossovers, some regions of local high crossover frequency remain, potentially due to introgression from the DH inducer which has been previously observed (Li et al. 2009) or possible factors such as residual heterozygosity in founder inbreds.

## QTL Mapping

### QTL Mapping and SNP Association

To analyze flowering time and plant height, we took both a linkage mapping and SNP association approach. Linkage mapping relies on linear regression of the phenotypes on the matrix of founder probabilities, while SNP association regresses the phenotypes on the biallelic marker states. We found high concordance between linkage mapping and association analysis, where the most significant loci were identified for all traits by both approaches (Figures S3 and S4). Thus, we will refer to the linkage mapping results to represent our findings.

### Flowering Time

Mapping for AnthGDU revealed several significant peaks across the ten chromosomes in the WI-SS-MAGIC DH population (Figure 4A, Figure S5). The most significant peaks appear on chromosome three at 156.3 and 163.1 Mbp and chromosome eight at 127.9 Mbp (Table S7). Peaks for anthesis and silking highly colocalized, which is expected due to the high correlation of the phenotypic values at r=0.83 (Figure S3). Several of the significant loci are near the location of other known flowering time genes. For example, chromosome three contains *ZmMADS69*, also known as *Zmm22*, and chromosome eight contains *ZmRap2.7* and *ZCN8*, as noted on Figure 4A. All three genes are known to regulate flowering time (Guo et al. 2018; Liang et al. 2019). Despite the high correlations between *per se* and hybrid flowering, several QTL that are significant in the *per se* population lose their significance in one or both hybrid populations. The large peak on chromosome eight containing *ZmRap2.7* and *ZCN8* is not significant in the SS-PHJ89 population, despite retaining its significance in the group of DH lines used to make the hybrid population (Figure 4C). The same locus remains significant in the SS-3IIH6 population, albeit with a smaller LOD score (Figure 4B). Loss of significance indicates a loss in variation, such that the tester may have a dominant allele that masks the variation within the DH population. This suggests that there are contrasting loci present between the two testers, where PHJ89 has a dominant locus relative to the *per se* population while 3IIH6 does not.

**Figure 4:**
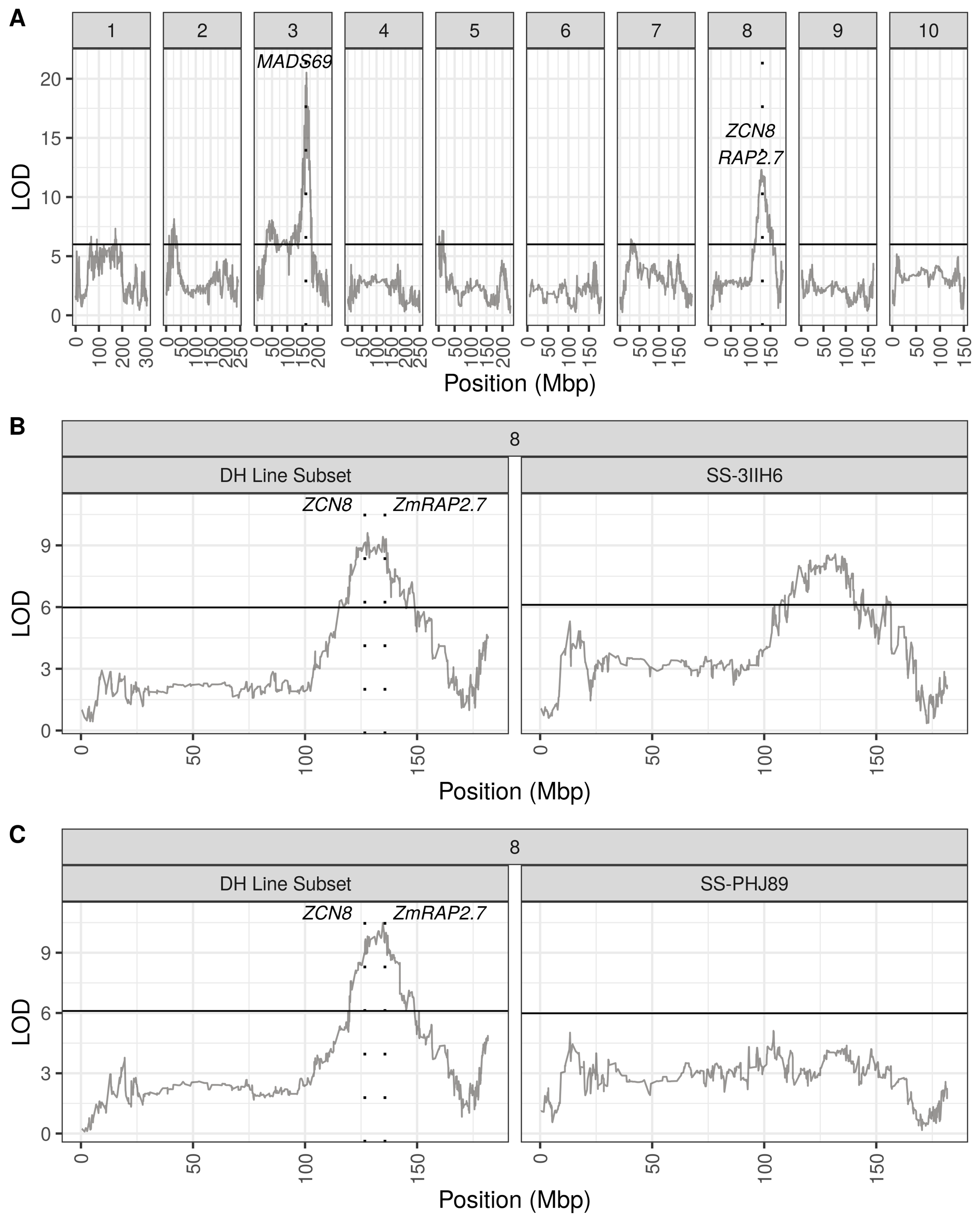
Anthesis GDU QTL mapping Population specific LOD scores are plotted for each panel. Dashed vertical lines show known flowering time genes. (A) QTL peaks for the *per se* population for each of the ten chromosomes. (B) QTL peaks for chromosome eight of the SS-3IIH6 population and the DH lines that were used to generate the population. (C) QTL peaks for chromosome eight of the SS-PHJ89 population and the DH lines that were used to generate the population.

We calculated BLUP QTL effects for *per se* AnthGDU on chromosomes three and eight and found allelic series at the significant loci on both chromosomes (Figure 5). On chromosome three, PHB47 provides the early flowering allele and LH145 provides the late flowering allele, while on chromosome eight PHJ40 provides the early allele, and B73 and B84 provide later alleles. It is notable that LH145 is the second earliest parent of the population and PHB47 flowers near the mean of the population, demonstrating that alleles for early and late flowering segregate within the parents. Using a single QTL model to fit the BLUP effects for the chromosome three peak at 163,105,981 bp, the most extreme alleles from the parents show a -27.2 +/- 8.8 GDU effect for PHB47 and +22.6 +/- 8.9 GDU effect for LH145. For the peak on chromosome eight at 127,898,534 bp, the most extreme effects are -24.5 +/- 8.9 GDU from PHJ40 and +17.1 +/- 8.9 GDU from B73.

**Figure 5:**
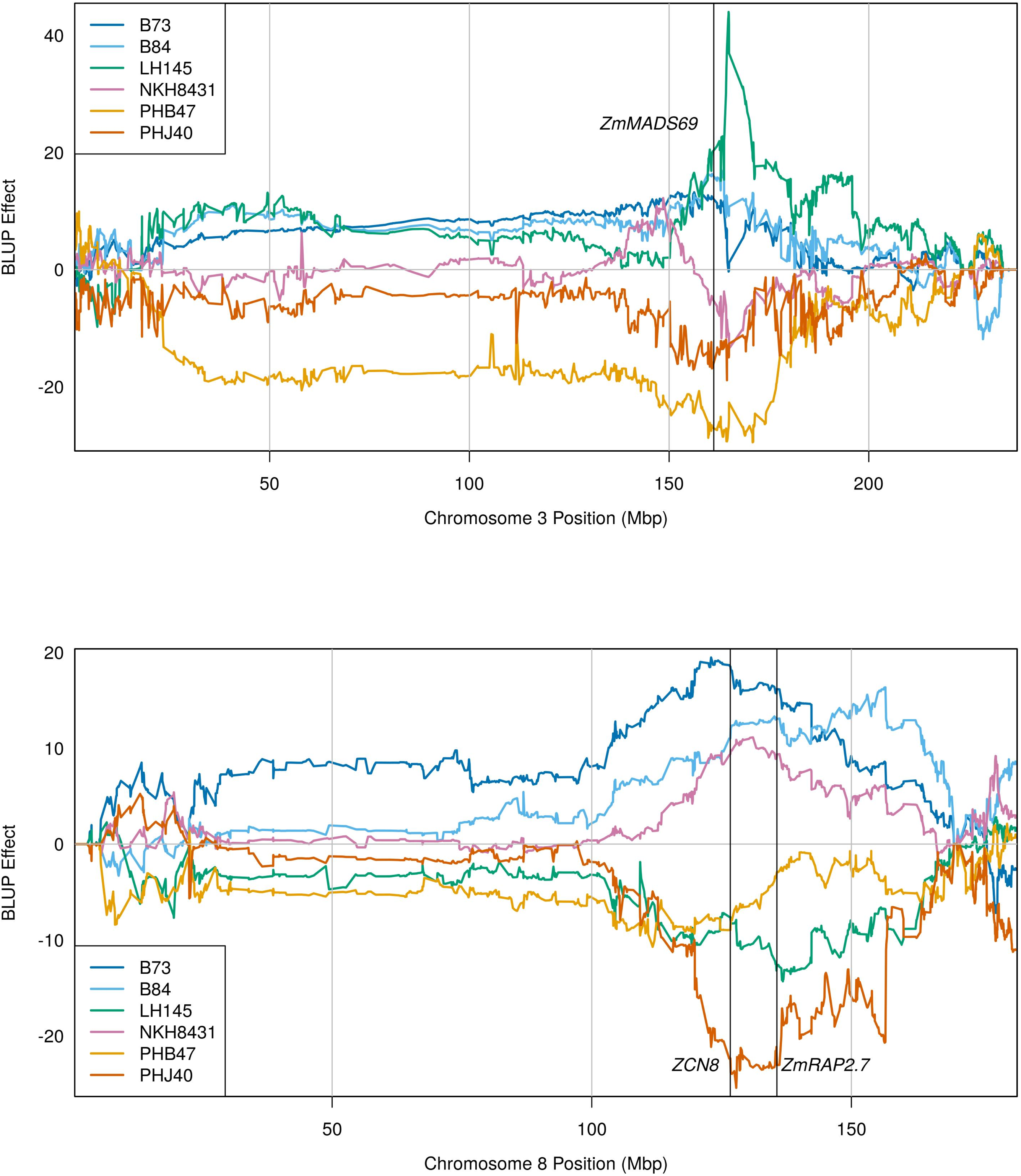
Founder anthesis GDU QTL BLUP effects BLUP effects for each parental contribution are plotted for the two chromosomes containing the most significant peaks for flowering time. Vertical lines are plotted denoting three major flowering time loci.

### Plant Height

Like flowering time, many significant peaks were also found for plant and ear height, such as on chromosomes one, two, three, and ten (Figure 6A, Figure S5, and Table S7). The most significant locus on chromosome one is located at 225.4 Mbp and it contains *brd1*, a gene which is involved in the brassinosteroid pathway and for which a mutant allele causes severe dwarfism (Makarevitch et al. 2012). Likewise, the most significant locus on chromosome three at 8.9 Mbp contains another gene discovered through mutational studies, *DWARF1* (*d1*), which is involved in the gibberellin pathway (Chen et al. 2014). Fewer loci colocalized for plant and ear height compared to AnthGDU and SilkGDU (Figure S4). Most interestingly, a peak on chromosome six at 105.8 Mbp appears for plant height in the SS-3IIH6 population, but not in the SS-PHJ89 population (Figure 5B). This peak is near *ubi3*, previously found to be associated with height traits (Ding et al. 2016). Previous studies have identified an epistatic interaction between *ubi3* and *br2* (Xiao et al. 2021), so we investigated the parental effects of the peak on chromosome six at 105,826,214 bp for both hybrid populations and found that the LH145 allele had an effect of +3.5 +/- 1.7 cm, while the B73 allele had an effect of -4.7 +/- 2.0 cm in the SS-3IIH6 population (Figure S6). Here again, *per se* B73 is the tallest of the parents while *per se* LH145 is the second shortest and their allelic effects are opposite of their overall phenotypes, but their hybrid phenotypes are both closer to the population mean. For comparison, the insignificant chromosome six locus in the *per se* and SS-PHJ89 populations shows no such differentiation between the parents (Figure S5). The genetic variance for plant height in the SS-3IIH6 population was 151.1 cm^2^, so the allelic effects here are a small proportion of the total variance.

**Figure 6:**
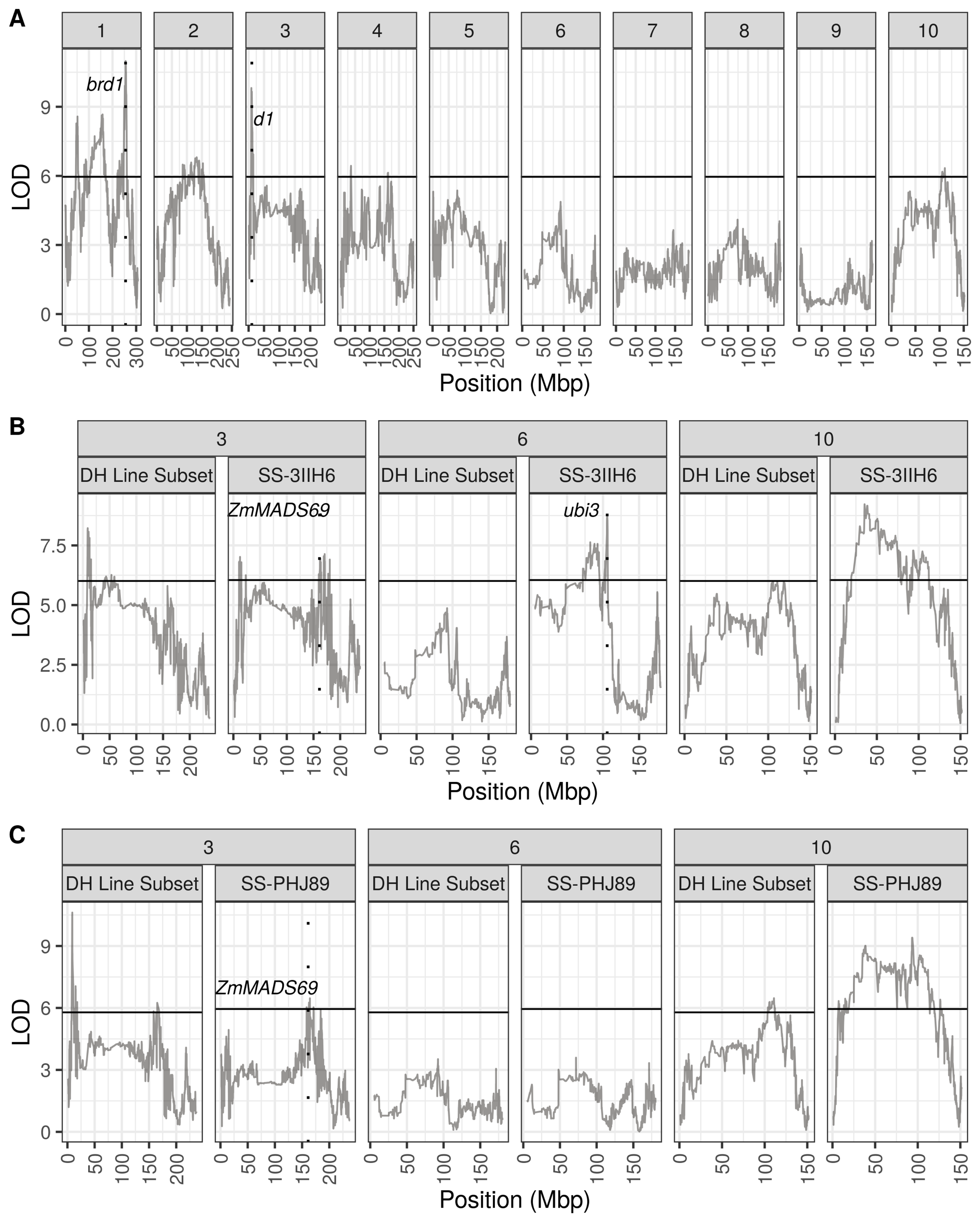
Plant height QTL mapping Population specific LOD scores are plotted for each panel. Dashed vertical lines show known height genes. (A) QTL peaks for the *per se* population for each of the ten chromosomes. (B) QTL peaks for three chromosomes of the SS-3IIH6 population and the DH lines that were used to generate the population. (C) QTL peaks for three chromosomes of the SS-PHJ89 population and the DH lines that were used to generate the population.

### Genomic Prediction

Because information on the DH lines was available before hybrid test crosses were made, we tested the predictive abilities of several direct and indirect genomic prediction models (Figure 7). As expected, the most successful models were those that were trained on the data that was most directly related to the predicted set, such as prediction within the hybrid SS-PHJ89 set and within the SS-3IIH6 set for plant height (r=0.63 and r=0.60, respectively) and prediction within the *per se* set for anthesis (r=0.61). Predictive abilities between test cross populations were moderate, with AnthGDU predictive abilities for SS-3IIH6 to SS-PHJ89 and vice versa of r=0.55 and 0.54, and plant height at 0.57 and 0.53, respectively. The correlations of the indirect predictions of *per se* to hybrid phenotypes were lower, but still greater than r=0.48. It is important to note that correlations do not consider the difference in scale between the *per se* and hybrid populations and cannot account for the population mean heterotic effect on both flowering time and plant height between the populations. Finally, we wanted to test the feasibility of using predicted *per se* data to discard DH lines from our breeding program that either are too tall or flower too late for our environment. We compared the predicted *per se* AnthGDU and *per se* plant height values to their observed values, and colored DH lines based on their status in the top 15^th^ percentile for either the predicted or observed value (Figure S7). We maintained this color scheme when plotting the DH line’s observed hybrid values to assess the combination of genomic prediction ability and tester response. Overall, we found that the DH lines in the top 15^th^ percentile for the predicted trait but not for the observed trait tended to be the DH lines that would make hybrids that are satisfactory to our breeding program’s needs, while DH lines that were in the top 15^th^ percentile of observed values tended to have higher hybrid values. These results are expected, especially considering the high correlations between *per se* and hybrid phenotypes and the lower predictive ability of the *per se* to hybrid models.

**Figure 7:**
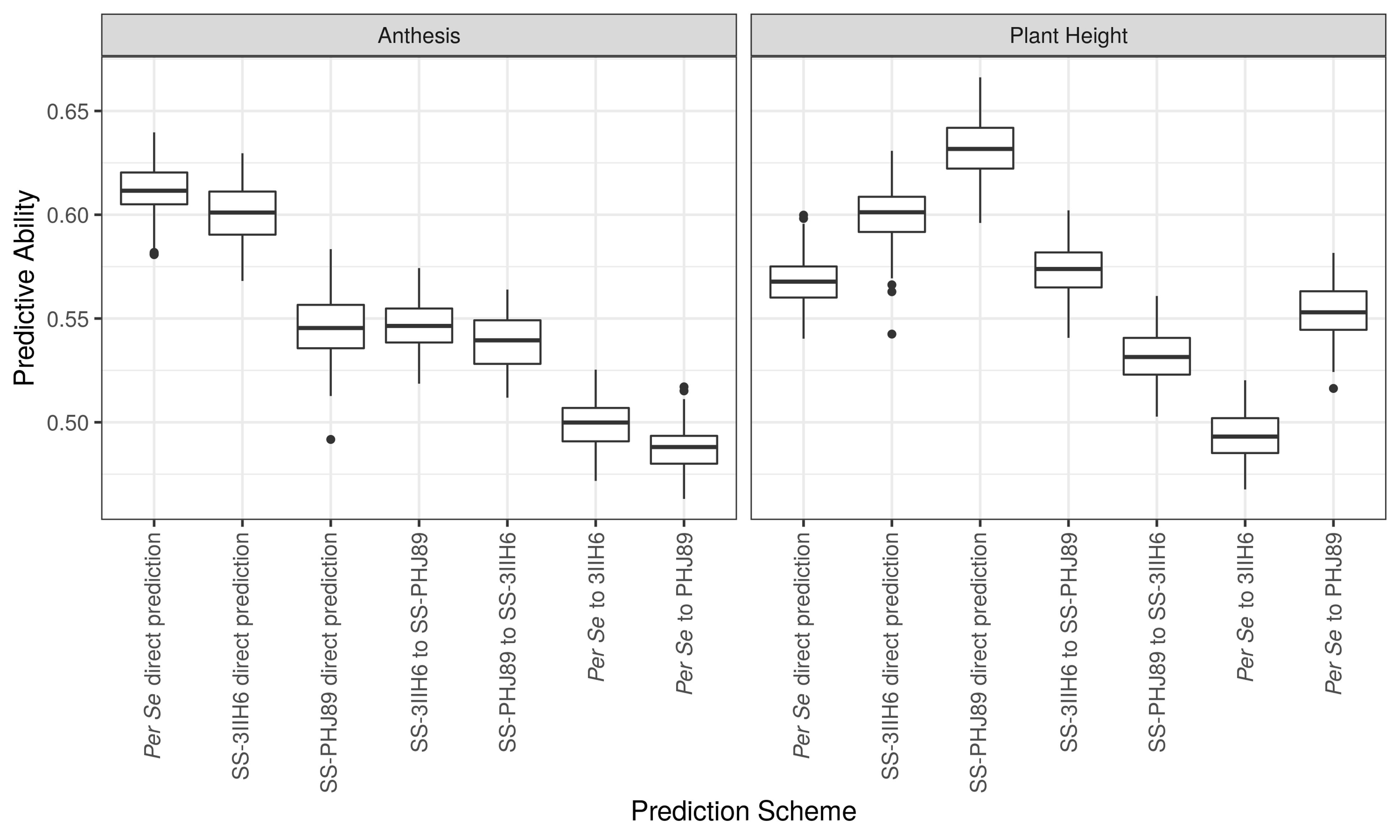
Predictive abilities for direct and indirect genomic prediction models Legend: Predictive abilities for 100 replications of each model. Direct models were trained using the population phenotype they would predict, while indirect models were trained with the *per se* or opposite hybrid population and used to predict each phenotype.

## DISCUSSION

### QTL mapping in multi-parent populations

Several multi-parent populations have been developed in maize, including MAGIC populations from Italy and Spain (Dell’Acqua et al. 2015; Jiménez-Galindo et al. 2019), four-parent populations from China and the US (Ding et al. 2015; Mahan et al. 2018), and nested association mapping populations from the US, China, and Europe (Yu et al. 2008; Li et al. 2013; Bauer et al. 2013; Giraud et al. 2017). The existing MAGIC, US-NAM, and CN-NAM populations use a variety of inbreds that sample the diversity of maize genetics, and the European NAM populations focus on the Dent and Flint heterotic groups in addition to factorial crosses made between recombinant inbred lines. Our population concentrates the founders within the Stiff Stalk heterotic group. An advantage to focusing on the Stiff Stalk group is that maize breeding relies on recycling genetics within heterotic groups to make new parents and crossing parents between groups to make hybrids. In a factorial mating design between Flint and Dent multiparent populations, it was discovered that the majority of general combining ability QTL were specific to one heterotic group (Giraud et al. 2017; Seye et al. 2019). Thus, blending the genomes of parents within a single heterotic group versus across the diversity of maize creates a more applicable population to study the subset of alleles present within Stiff Stalk seed parent germplasm released in North America. Breeding based on heterotic groups is expected to drive diverging allele frequencies between groups and constraining our mapping population to a single heterotic group allows us to examine the effects of these alleles on agronomic and yield related traits within their intended context. Mixing multiple founders takes advantage of historical recombination in addition to recombination introduced through population development. Multiple founders within a single population allows the study of allelic series at loci of interest, such as for AnthGDU on chromosomes eight and three (Figure 5).

### QTL mapping in the WI-SS-MAGIC

Large efforts have been made by the plant research community to elucidate the control of complex traits such as flowering time and height. Our results confirm the previous findings of numerous authors, especially the candidate genes highlighted in Figures 4 and 6. Similar loci were found within the *per se* and hybrid populations, but there was variation between the populations, indicating that non-additive variation impacts the hybrid phenotypes (Supplemental Figure 5). For flowering time, the loss of significance of the QTL on chromosome eight indicates the PHJ89 tester has a completely dominant locus compared to the *per se* population and compared to the other tester, 3IIH6. Our results demonstrate that QTL detection depends on the genetics of the tester when mapping in hybrid populations. While it is possible that the absence of a signal in the hybrid population could be due to environmental or genotype by environment effects, the high heritabilities support the large role of genetic variation.

For plant height, a significant locus exists within the hybrid SS-3IIH6 population that is absent in the *per se* population, which could indicate an epistatic interaction between the tester and population genotypes. This locus has been previously described in the context of both inbred and hybrid populations (Ding et al. 2016; Xiao et al. 2021). Ding et al. (2016), used a near isogenic line from the US-NAM family B73 × Tzi8 crossed to B73 to finemap the QTL to 95-96 Mbp on chromosome six. Like our study, Xiao et al. (2021), found a plant height QTL near 95.8 Mbp on chromosome six within one test cross population but not in the inbred population, and provides a schematic outlining the epistatic derepression uncovered by this locus. In theory, there is the potential to study epistasis between more than two founders within a MAGIC population. For the WI-SS-MAGIC population, comparing the founder states for two loci results in 36 total digenic classes. We separated our population into these classes for the two most significant loci for *per se* AnthGDU and found the mean number of individuals per class was 16.5, with a range of seven to 44 individuals (Supplementary Figure 8). Smaller population size in some of the sets exacerbates this issue of power. We repeated the procedure for the SS-3IIH6 plant height loci on chromosome one and six and found a mean of 11.0 individuals per class, with a range of zero to 27, and nine classes have fewer than five observed individuals (Supplementary Figure 9). The limited number of observations per digenic class restricts the ability to statistically evaluate interaction between loci.

### QTL mapping in DH lines and hybrids

Previous work in mapping QTL across testers has found high concordance between plant height QTL discovered in different test cross populations and minimal digenic epistasis, despite evidence for epistasis under generation means analysis (Lübberstedt et al. 1997; Melchinger et al. 1998). Tester relatedness also influences the ability to discover QTL, where a tester unrelated to the population uncovers QTL for additive traits more effectively than related testers (Frascaroli et al. 2009). Another study of a biparental RIL population crossed to four testers found that mapping the mean test cross height was sufficient to identify shared loci between testers (Austin et al. 2001). Recent work by Xiao et al. (2021), examining heterosis for over 42,000 hybrids generated by crossing 1428 multi-parent lines with 30 testers found that epistasis plays a role in generating heterosis, contradicting previous work demonstrating the low impact of epistasis (Hinze and Lamkey 2003; Mihaljevic et al. 2005).

We documented our population’s response to two testers to better understand the heterotic effect of Iodent (3IIH6) and Oh43 (PHJ89) type testers on the WI-SS-MAGIC population. Understanding *per se* phenotypes requires mapping in an inbred population, while understanding an inbred’s response to a tester requires evaluating and mapping traits in the hybrid population. We found the hybrid populations to have less than half the variation of the *per se* population, which could indicate that non-additive gene action is affecting the phenotypes. The interplay of dominant and recessive loci only manifests in the hybrid population, either creating or concealing phenotypic variation depending on the gene action of the trait. Our study found evidence for contrasting allelic states between the two testers in several regions of the genome based on disappearance and appearance of QTL, including a dominant locus for flowering time on chromosome eight in PHJ89 compared to 3IIH6, and a putatively epistatic locus revealed for plant height in 3IIH6. Despite the strong positive correlation between the hybrid phenotypes, several loci were found in only one of the hybrid populations (Supplementary Figure 5). Perfect correlation between the test cross phenotypes would lead to the discovery of the same QTL between the two populations, yet the deviation from a one-to-one relationship between the test cross phenotypes leads to the discovery of unique QTL in the hybrid populations. Choice of tester influences hybrid performance and QTL mapping results, as evidenced by studies previously described and our findings. Despite the high correlation between the phenotypes of the hybrid populations and the expected identity by descent between the two testers, unique QTL were discovered for each trait in each population.

### Genomic prediction of hybrid phenotypes

In maize breeding programs, *per se* phenotypes are often available before hybrid varieties can be tested. Using *per se* and hybrid data from our study, we investigated the association between *per se* and hybrid flowering and height traits. We wanted to test the feasibility of predicting correlations of hybrid flowering time and height based on DH line measurements for the purposes of discarding lines that flower too late or are too tall for our breeding program. We also wanted to evaluate the predictive ability between the two hybrid populations as breeding programs often use multiple testers as materials advance through selection pipelines.

We found that the hybrids flowered earlier and were taller than their maternal DH parents, confirming heterotic relationships for flowering and height found in other studies (Flint-Garcia et al. 2009; Li et al. 2018). Previous studies have found increased predictive abilities when incorporating parental inbred information (Liang et al. 2018; Jarquin et al. 2020), and we also found moderate prediction abilities for hybrid flowering time and plant height when the models were trained using the *per se* data. As expected, the models with the highest predictive abilities were those that were trained on the data they were designed to predict, although we achieved predictive abilities between r=0.49 and r=0.55 for models predicting hybrid traits that were trained with *per se* data.

Correlations between *per se* and hybrid populations do not consider the difference in magnitude between them, such as the average difference between *per se* and hybrid flowering of 90 GDU or difference in height of 71 cm. Heterosis due to small genome-wide effects produces a relatively uniform incremental decrease in flowering time and increase in height across all lines, while variation within inbreds and within hybrids is largely due to similar large effect QTL likely in combination with undetected small effect loci. These findings are consistent with the overlapping and non-overlapping QTL that were found between the *per se* and hybrid populations because the difference between the predictive abilities for direct and *per se* to hybrid models cannot account for the dominance or epistatic effect of the tester at individual loci. In addition, the masking of *per se* QTLs within either of the hybrid populations is conceptually consistent with the lowered predictive ability of using *per se* data to predict hybrids. We also found that the errors between predicted and observed *per se* phenotypes were a source of selection error that led to discarding DH lines that would have generated acceptable hybrids.

Finally, we also used the highest associated SNP from each LOD peak as fixed effects in the genomic prediction model but found that including the fixed marker effects lowered the predictive ability compared to using only the realized relationship matrix (data not shown). This finding supports previously simulated results demonstrating that known genes are only beneficial to models when they are few in number and explain large proportions of the variance (Bernardo 2014).

### Relevance to maize breeding

This method has applications in maize breeding because genomic prediction could be used to make selections prior to generating test cross seed for an entire population. Alternatively, a smaller subset of an inbred population could be grown as a model training set with several testers prior to generating larger hybrid populations. Genomic prediction could then be used to discard the poorest performing lines, which would increase genetic gain by increasing the selection intensity on the population. Overall, our results indicate that plant breeders should be less aggressive when using predicted *per se* data to predict hybrid performance because the errors in genomic prediction can lead to discarding hybrid lines incorrectly. Plant breeders must balance their selection for maize yield with the adaptation requirements and architectural risk for root or stalk lodging when developing new inbred lines, as demonstrated by including information for flowering time and plant height in this study. Flowering time and moisture at harvest are also indications of overall relative maturity, which is an important characteristic that plant breeders use to place varieties across geographies and that farmers use to balance risk and make planting time and cultivar decisions. Our results indicate that flowering time and height have high correlations between DH lines and hybrids within these DH line-tester combinations yet experience different profiles of QTL significance across the genome. While the goal of maize breeding efforts is to increase or protect hybrid yield, most genetic research efforts focus on using inbreds to study complex traits. Understanding how traits manifest in a parental inbred versus its hybrid progeny is a critical area of maize breeding and quantitative genetics research. For example, parental *per se* measures of grain yield have been previously used to increase prediction ability of hybrid performance (Schrag et al. 2010). Deviations from the purely additive relationship of inbred flowering time or plant height to hybrid phenotype can be investigated to add to the underlying understanding of the gene action that supports genomic prediction.

## CONCLUSIONS

In conclusion, several known loci were uncovered in different combinations within the *per se* and test cross sets of a MAGIC population. Dominance of one tester over the population caused the loss of a highly significant peak for anthesis, while the presence of the other tester revealed putative epistatic variation for plant height. The six parents of the population are all members of the Stiff Stalk heterotic group, which is the canonical source of seed parent germplasm in the United States. These lines represent the diversity of both the sub-heterotic groups within the Stiff Stalk pool and the major plant breeding entities operating in the 1970’s and 1980’s and are no longer under intellectual property protection. In addition to dissecting the genetic architecture of these complex traits, this study provides a description of a new population resource available to maize researchers. Multi-parent populations are a unique mapping resource to study the effect of more than two parental alleles on quantitative traits, and they are a means to increase the diversity of alleles under study while managing minor allele frequency.

Further, the genome assemblies of the six parents with annotation from a five-tissue transcriptome atlas (Li et al. 2021; Bornowski et al. 2021) are available for study, which increases the variety of opportunities for maize researchers. This population could be used to assay the effect of the alleles present within the population on combining ability, adaptation, genotype by environment interaction, stability, and provide a new paradigm for studying traditional and genomic selection. The practicality of leveraging linkage mapping of highly polygenic traits to make selections within breeding programs has been limited in the past, especially for traits that follow an infinitesimal model such as maize height (Peiffer et al. 2014). Nevertheless, further study of individual loci can impact plant breeding through mutational studies made possible by gene editing, in addition to current efforts in commercial plant breeding accomplished through genomic selection.

## Data Availability

All supplementary tables and files have been uploaded to FigShare. Supplemental file S1 contains summary statistics regarding the exome capture, supplemental file S2 contains the exome capture probe coordinates, supplemental file S3 contains supplemental methods, supplemental file S4 is contains the unfiltered 1.8 million SNPs data, and supplemental file S5 contains the control folder for mapping in R/qtl2, including marker maps, genotypes, phenotypes, and cross information. Supplemental table ST1 contains descriptions of the lines in the study, ST2 contains details about the field experiments, ST3 contains plot-based data, ST4 contains information about the stage one models used for calculating BLUE phenotypes, ST5 contains the BLUE phenotypes, ST6 contains the reference range haplotype IDs, and ST7 contains all significant QTL peaks. Raw exome capture and GBS sequence reads are available in the National Center for Biotechnology Information Short Read Archive under BioProjects PRJNA781987 and PRJNA781986 respectively.

## Acknowledgements

The authors acknowledge Mona Mazaheri and Alden Perkins for support with tissue sampling, Brieanne Vaillancourt and Joshua C. Wood for assistance with exome capture sequence data processing, Zachary Miller, Lynn Johnson, Joseph Gage for advice in building the WI-SS-MAGIC Practical Haplotype Graph, Dylan Schoemaker for helpful discussion on genomic prediction and the manuscript, and AgReliant Genetics for providing doubled haploid induction as in-kind support. The authors utilized the University of Wisconsin – Madison Biotechnology Center’s DNA Sequencing Facility (Research Resource Identifier – RRID:SCR_017759) to extract DNA, generate GBS libraries, and sequence GBS libraries. The UWBC is a Licensed Service Provider for internal and external clients, providing GBS services under license from Keygene N.V. which owns patents and patent applications protecting its Sequence Based Genotyping technologies.

## Funding

This work was supported and funded by the U.S. Department of Energy Great Lakes Bioenergy Research Center (DOE BER Office of Science DE-FC02-07ER64494) to CRB, SMK, NdL; USDOE ARPA-E ROOTS Award Number DE-AR0000821, National Corn Growers Association and Iowa Corn Growers Association support to DCL, NdL, SMK; the D.C. Smith Wisconsin Distinguished Graduate Fellowship to KJM; the National Institute of Food and Agriculture, United States Department of Agriculture Hatch 1013139 and 1022702 project to SMK, and National Institutes of Health grant R01GM070683 to KWB. The work (proposal: 10.46936/10.25585/60000838) conducted by the U.S. Department of Energy Joint Genome Institute (https://ror.org/04xm1d337), a DOE Office of Science User Facility, is supported by the Office of Science of the U.S. Department of Energy operated under Contract No. DE-AC02-05CH11231.

## Conflict of interest

The authors declare no conflicts of interest.

## Abbreviations

BSSS: Iowa Stiff Stalk Synthetic
BLUE: best linear unbiased estimator
DH: doubled haploid
ex-PVP: expired Plant Variety Protection
GWAS: genome wide association study
MAGIC: multi-parent advanced generation intercross population
PHG: Practical Haplotype Graph
PHI: Pioneer Hi-Bred, International
PVP: Plant Variety Protection
QTL: quantitative trait locus or loci
SS: Stiff Stalk

---

**Supplemental Figure 1:**
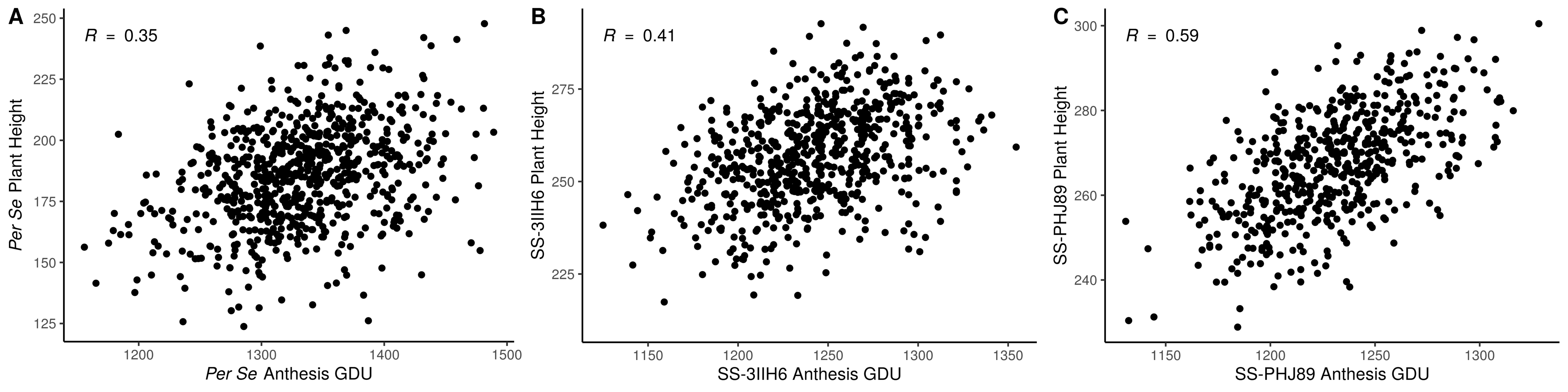
Phenotypic correlations between traits Scatterplots of BLUEs demonstrate the positive correlation between traits, within populations. Pearson correlations are shown in the upper left.

**Supplemental Figure 2:**
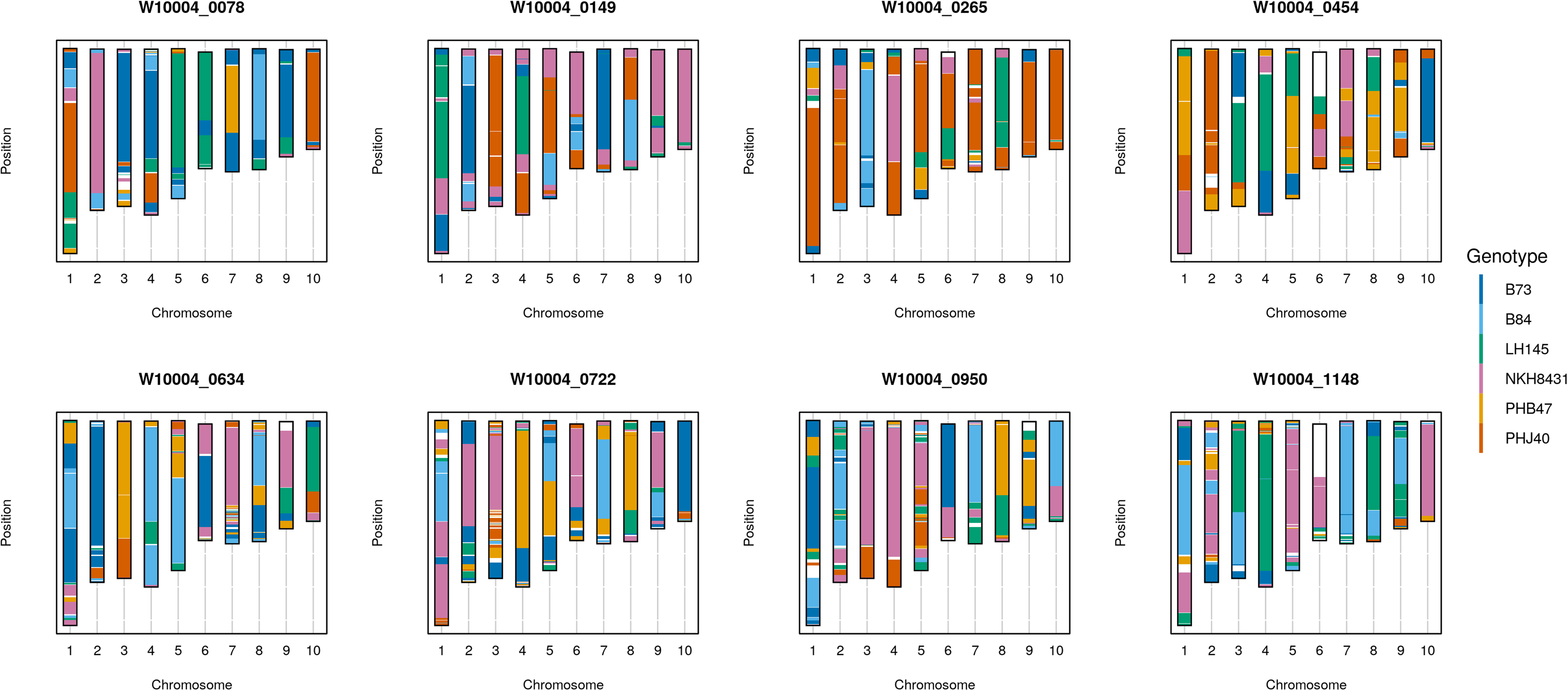
Haplotypes for six representative UW-SS-MAGIC lines Lines in the top row are from Subset A, and each have 60 crossovers. Lines in the bottom row are from Subset B, and each have 103 crossovers. Genomic areas plotted in white did not have a founder probability rise above 50% in that region.

**Supplemental Figure 3:**
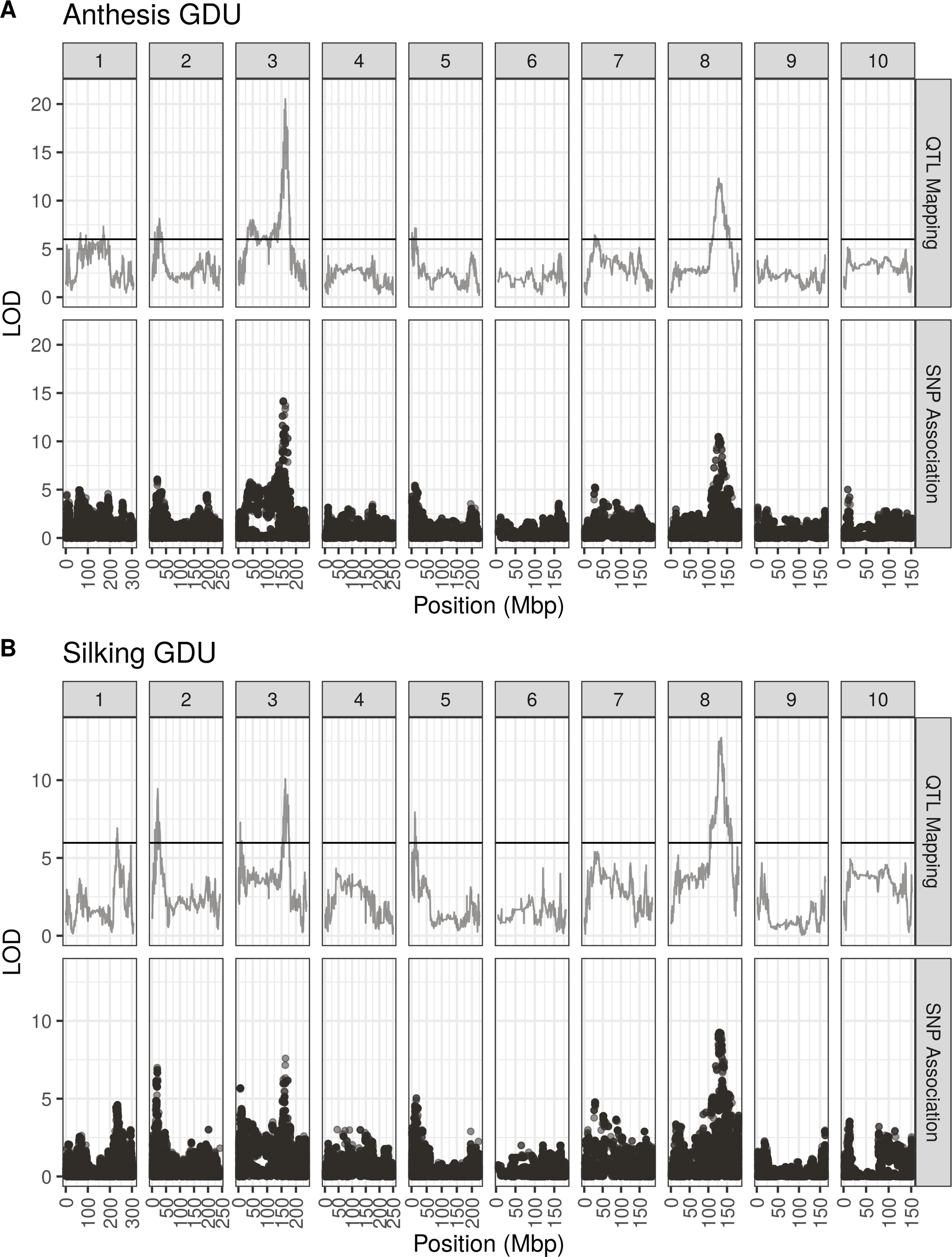
QTL linkage mapping and GWAS for flowering time (A) Anthesis GDU QTL peaks and SNP association results in the *per se* population. (B) Silking GDU QTL peaks and SNP association results in the *per se* population.

**Supplemental Figure 4:**
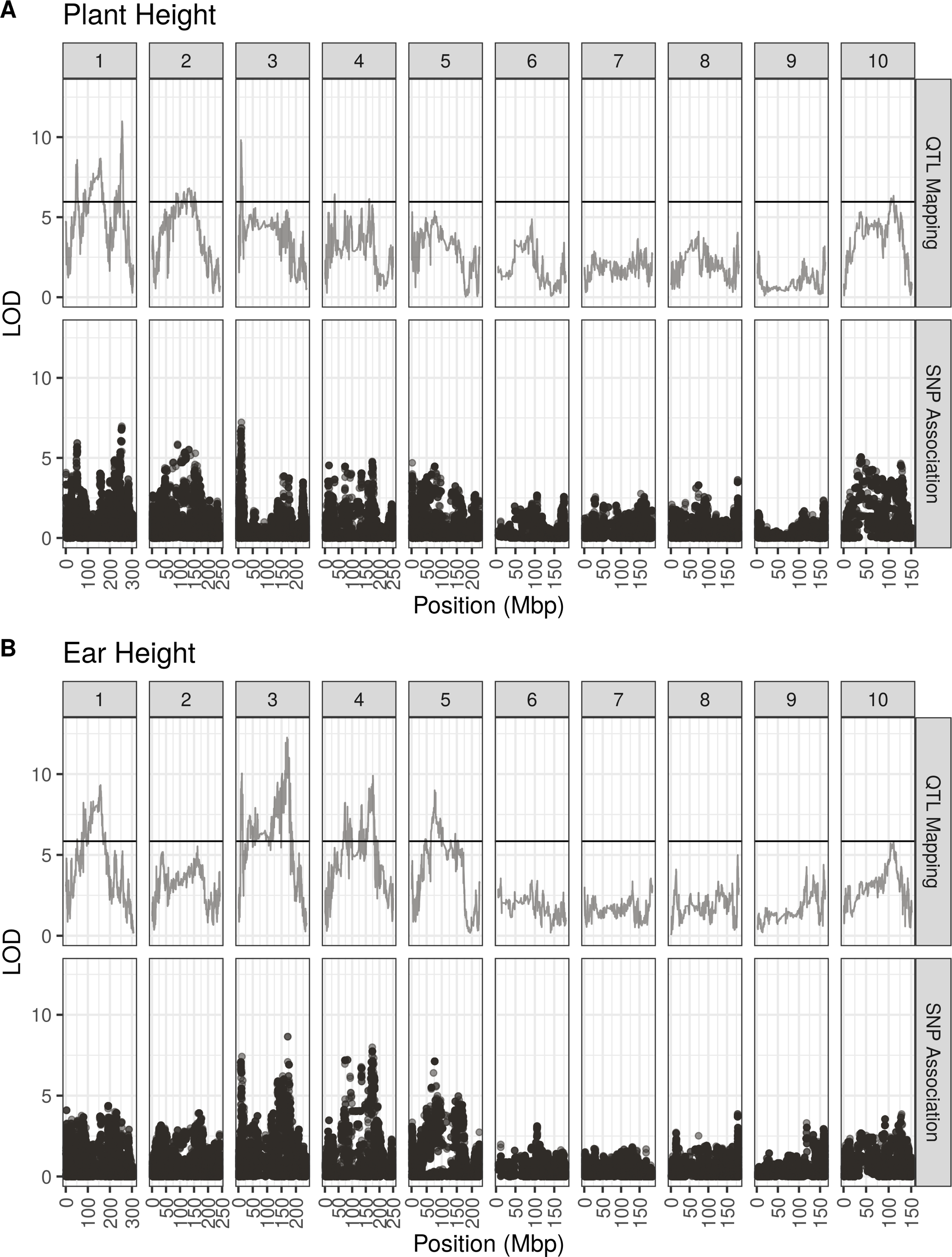
QTL linkage mapping and GWAS for plant and ear height (A) Plant height QTL peaks and SNP association results in the *per se* population. (B) Ear height QTL peaks and SNP association results in the *per se* population.

**Supplemental Figure 5:**
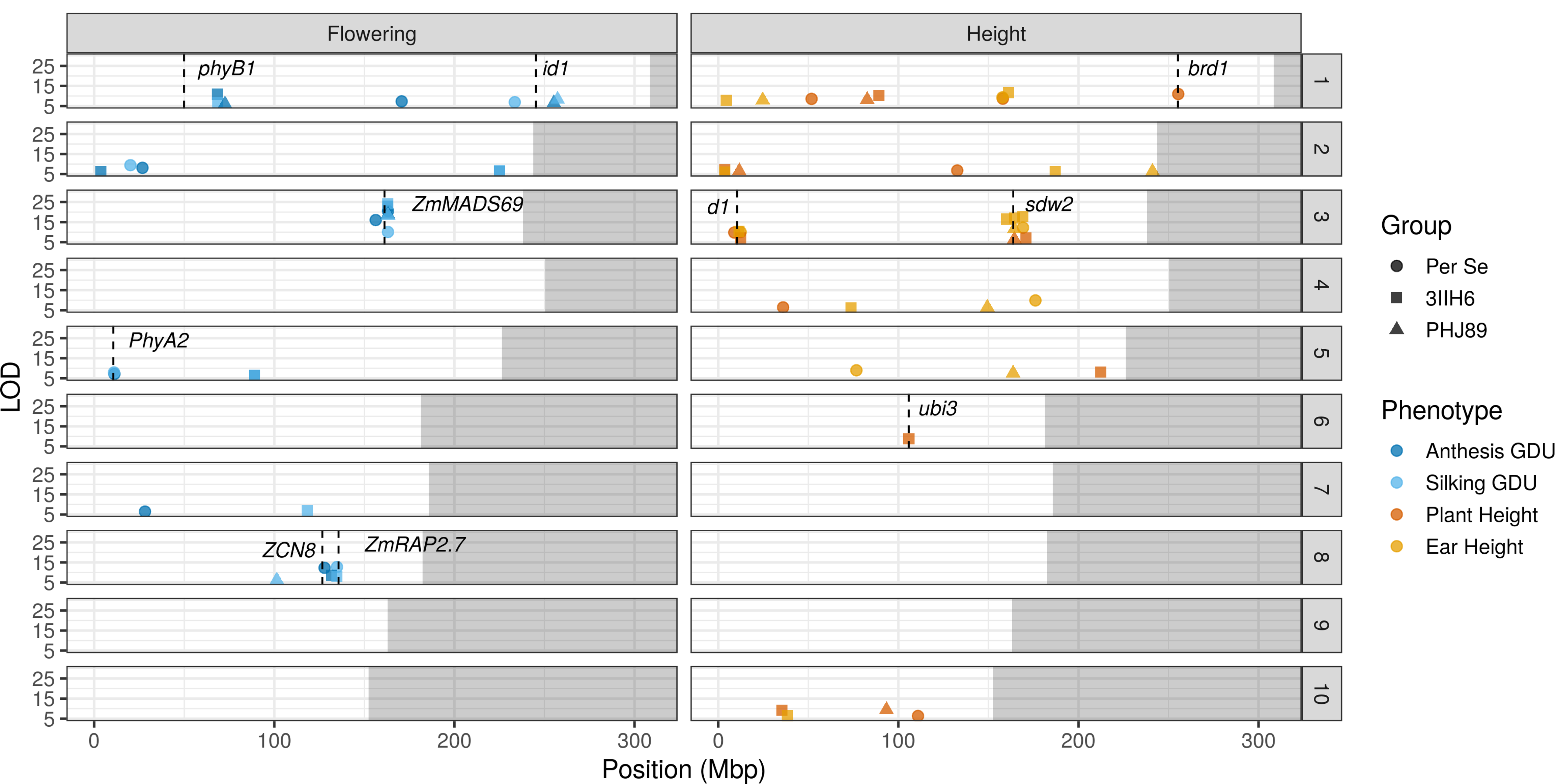
All significant QTL identified for flowering time and plant height Significant loci are plotted for flowering and height traits for the *per se* population, SS-3IIH6 hybrid population, and SS-PHJ89 hybrid populations. Previously characterized flowering and height loci are plotted as dashed vertical lines. To declare two significant loci under one large QTL peak, the LOD score was required to drop by at least five.

**Supplemental Figure 6:**
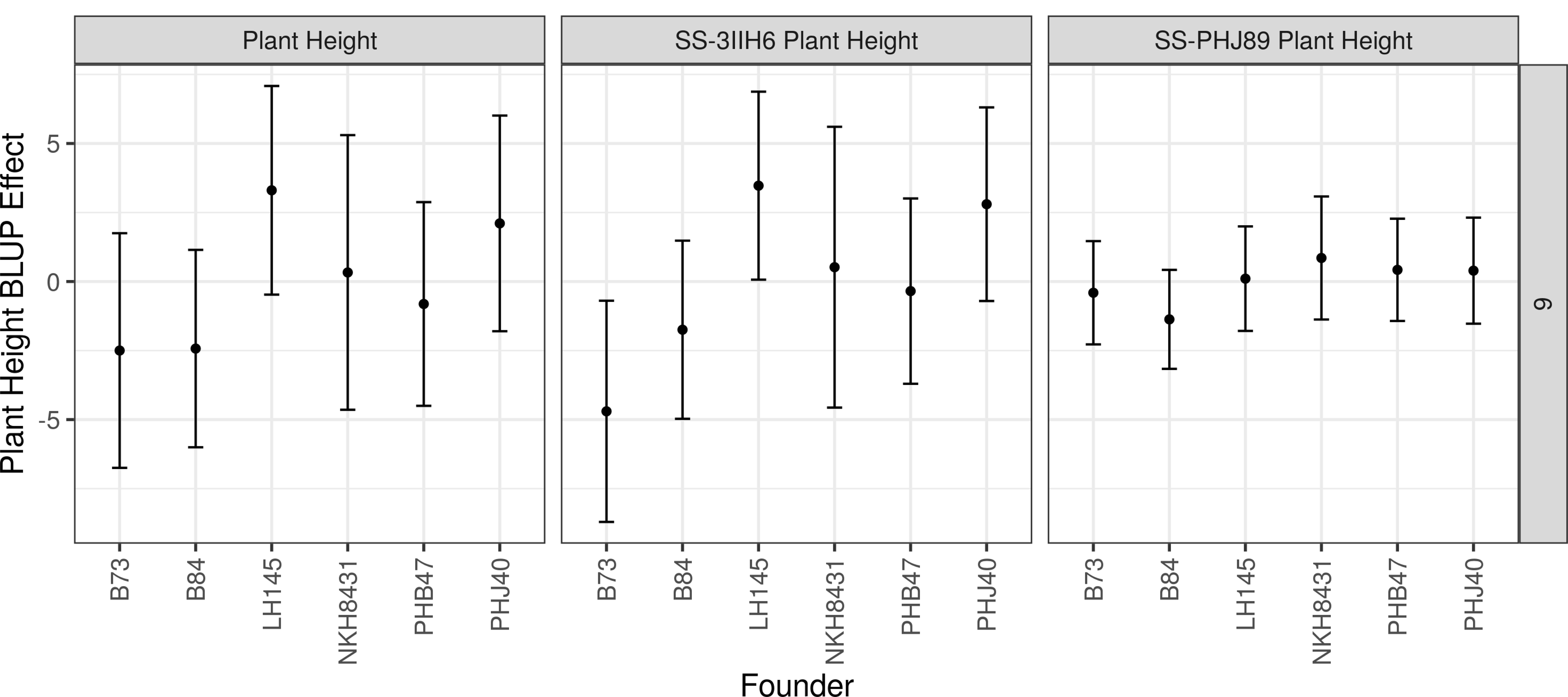
Founder plant height BLUP effects QTL BLUP effects with +/- 2 standard errors at the most significant locus for SS-3IIH6 plant height in the *per se*, SS-3IIH6, and SS-PHJ89 populations on chromosomes six. This QTL was not significant in the *per se* or SS-PHJ89 populations.

**Supplemental Figure 7:**
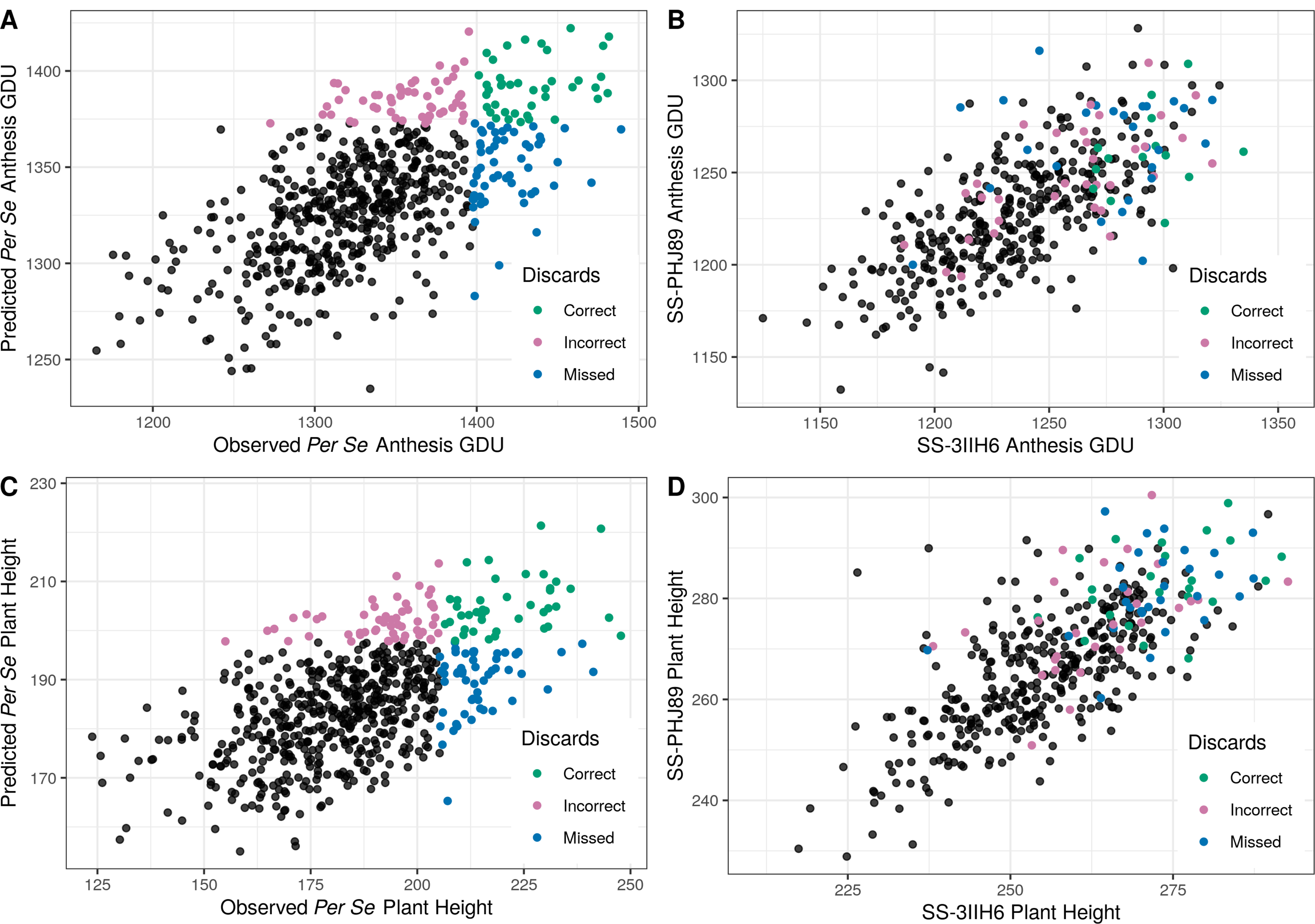
Observed vs predicted phenotypes and discard accuracy (A) Scatterplot of observed vs predicted anthesis GDU for the *per se* population. To select for adaptation to Wisconsin, our breeding program discards the latest lines of a population. Lines in the top 15% of both observed and predicted anthesis values (i.e. flower the latest) are colored in green, lines that flower in the latest 15% of predicted values are plotted in pink, and lines that flower in the latest 15% of observed values are plotted in blue. Color is recorded for each DH line name. (B) Using the DH line color scheme from A, the hybrid phenotypes are plotted for SS-3IIH6 and SS-PHJ89. (C) Scatterplot of observed vs predicted plant height for the *per se* population. Our breeding program discards the tallest members of the population. Lines are colored in the same manner as Panel A. (D) Using the DH line color scheme from C, the hybrid phenotypes are plotted for SS-3IIH6 and SS-PHJ89.

**Supplemental Figure 8:**
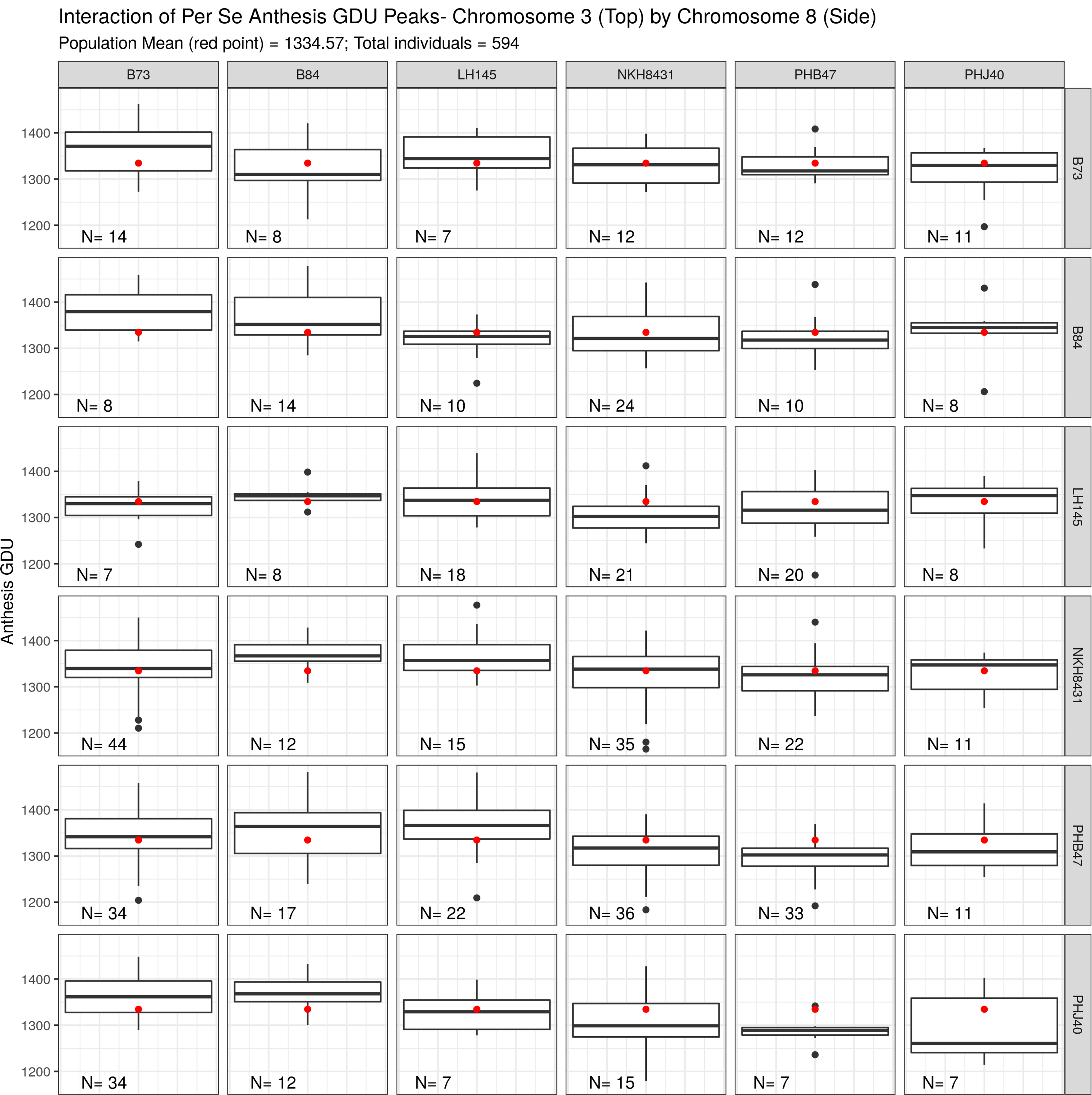
Box plots for each *per se* digenic class for two flowering time loci The study of epistasis is difficult due to low sample sizes of each digenic class. Each panel represents a digenic class for the two most significant *per se* anthesis GDU loci on chromosomes three and eight. The population wide mean is plotted as a red dot on each panel. Sample size for each class is in the lower left corner.

**Supplemental Figure 9:**
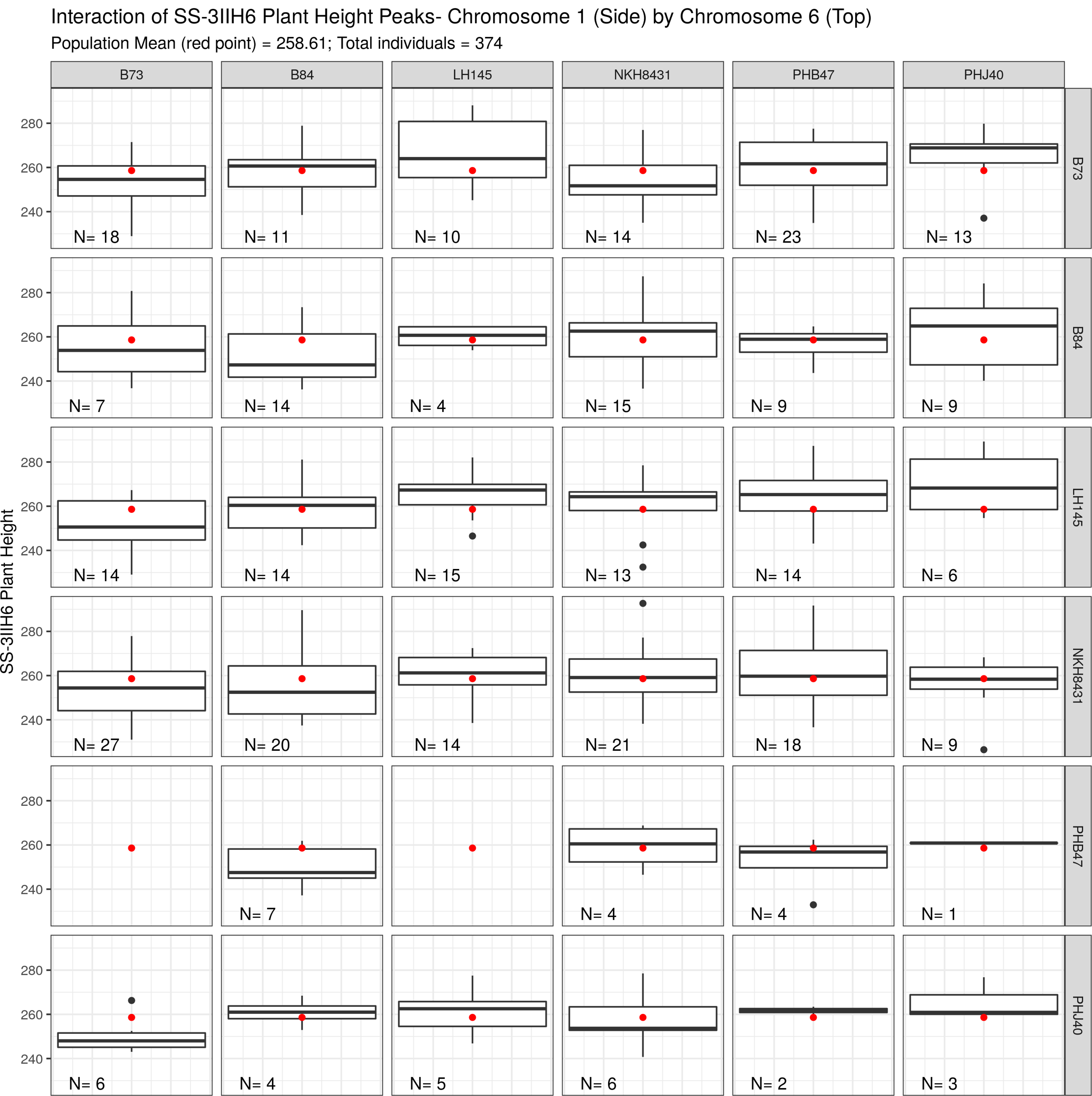
Bar plots for each SS-3IIH6 digenic class for two plant height loci The study of epistasis is difficult due to low sample sizes of each digenic class. Each panel represents a digenic class for two significant SS-3IIH6 plant height loci on chromosomes one and six. The population wide mean is plotted as a red dot on each panel. Sample size for each class is in the lower left corner. Some digenic classes have no observed individuals.

